# cGLRs are a diverse family of pattern recognition receptors in animal innate immunity

**DOI:** 10.1101/2023.02.22.529553

**Authors:** Yao Li, Kailey M. Slavik, Benjamin R. Morehouse, Carina C. de Oliveira Mann, Kepler Mears, Jingjing Liu, Dmitry Kashin, Frank Schwede, Philip J. Kranzusch

## Abstract

cGAS (cyclic GMP-AMP synthase) is an enzyme in human cells that controls an immune response to cytosolic DNA. Upon binding DNA, cGAS synthesizes a nucleotide signal 2′3′-cGAMP that activates the protein STING and downstream immunity. Here we discover cGAS-like receptors (cGLRs) constitute a major family of pattern recognition receptors in animal innate immunity. Building on recent analysis in *Drosophila*, we use a bioinformatic approach to identify >3,000 cGLRs present in nearly all metazoan phyla. A forward biochemical screen of 140 animal cGLRs reveals a conserved mechanism of signaling including response to dsDNA and dsRNA ligands and synthesis of alternative nucleotide signals including isomers of cGAMP and cUMP-AMP. Using structural biology, we explain how synthesis of distinct nucleotide signals enables cells to control discrete cGLR-STING signaling pathways. Together our results reveal cGLRs as a widespread family of pattern recognition receptors and establish molecular rules that govern nucleotide signaling in animal immunity.

## Introduction

Innate immunity is one of the first lines of defense that protects multicellular organisms from pathogen infection. In animal cells, a suite of protein sensors named pattern recognition receptors (PRRs) sense microbial replication by directly recognizing pathogen associated molecular patterns (PAMPs). Following PAMP ligand recognition, PRRs activate downstream innate immune responses through highly conserved transcriptional signaling pathways, inflammatory responses and cytokine release, and autophagy or cell death pathways that clear pathogen infected cells (Li and Wu, 2021; Takeuchi and Akira, 2010). Many of the core components that control innate immune signaling are evolutionarily ancient and broadly conserved throughout the metazoan kingdom. Pioneering studies in the invertebrate fruit fly model system *Drosophila melanogaster* discovered Toll as a PRR activated in response to fungi and Gram-positive bacteria (Lemaitre et al., 1996). These results led to the later identification of Toll-like receptors as PRRs in human cells (Medzhitov et al., 1997; Poltorak et al., 1998), highlighting how discovery of PRR function in invertebrate metazoans provides a critical foundation to define the general principles that control immune activation in animals (Akira et al., 2006; Flajnik and Kasahara, 2010).

The best understood PRRs in animals are categorized into four major sub-families: Toll-like receptors (TLRs), NOD-like receptors (NLRs), RIG-I-like receptors (RLRs), and C-type lectin receptors (CLRs) (Li and Wu, 2021; Takeuchi and Akira, 2010). Typically, animal genomes encode multiple genes for each PRR sub-family resulting in expression of closely related receptors that exhibit distinct preferences for PAMP ligand recognition. For example, human cells encode 10 TLRs that recognize diverse PAMPs including bacterial and fungal cell wall components (TLR2, TLR4, TLR6), single- and double-stranded nucleic acid (TLR3, TLR7, TLR8, TLR9), and bacterial flagellin protein (TLR5) (Fitzgerald and Kagan, 2020). PRRs including TLRs and NLRs are conserved in most metazoans, but genetic radiation events are known to dramatically expand or contract the number of individual protein family members within each animal species (Leulier and Lemaitre, 2008; Zhang et al., 2010). For example, mammals typically encode ∼20 NLR genes, while the zebrafish *D. rerio* and sea urchin *S. purpuratus* genomes encode >200 NLR family-members (Li et al., 2017; Meunier and Broz, 2017; Zhang et al., 2010), suggesting that rapid evolution of PRR gene families enables animal lineages to specifically tailor pathogen innate immune responses.

In mammals, cGAS (cyclic GMP–AMP synthase) is a PRR that senses double-stranded DNA mislocalized in the cell cytosol (Sun et al., 2013). Following dsDNA recognition, cGAS synthesizes the nucleotide second messenger signal 2′3′-cGAMP that directly binds and activates the receptor protein STING (stimulator of interferon genes) to initiate IRF3- and NF-κB-dependent transcriptional responses (Ablasser and Chen, 2019). cGAS-STING signaling is evolutionarily ancient and originated in bacteria as a potent form of antiviral defense (Whiteley et al., 2019; Cohen et al., 2019; Morehouse et al., 2020). Bacteria encode thousands of cGAS/DncV-like Nucleotidyltransferase (CD-NTase) enzymes that control highly divergent anti-phage defense signaling pathways (Whiteley et al., 2019; Lowey et al., 2020; Ye et al., 2020). Recently, a complete animal cGAS-like signaling pathway was discovered in *Drosophila* where cGAS-like receptor 1 (cGLR1) functions as a PRR that senses double-stranded RNA and synthesizes 3′2′-cGAMP to activate STING and restrict viral replication (Slavik et al., 2021; Holleufer et al., 2021). Considering the enormous diversity of CD-NTase enzymes in bacterial anti-phage defense, the discovery of *Drosophila* cGLR1 suggests that cGAS homologs may have a widespread role in animal innate immunity.

Here we define cGAS-like receptors (cGLRs) as a major family of PRRs in animal innate immunity. We identify >3,000 cGLRs with predicted complete catalytic active sites including representatives in nearly all major animal phyla. Using a forward, biochemical screen, we reconstitute cGLR signaling from representatives across the protein family tree and discover 15 new active animal cGLRs. cGLRs in divergent animal genomes respond to the common PAMPs dsDNA and dsRNA and synthesize nucleotide second messenger immune signals including novel products that contain pyrimidine bases. We show how diversification of cGLR nucleotide second messengers and STING receptors enables animals to establish complex networks for pathogen detection. Together, our results define the molecular rules that control cGLR signaling in animal innate immunity and explain how radiation of an ancient cGAS-like nucleotide signaling domain enabled diversification of pathogen sensing.

## Results

### Discovery of diverse cGLRs in animal innate immunity

Human cGAS and *Drosophila* cGLR1 share only ∼25% sequence identity at the amino acid level, strongly suggesting that these enzymes comprise only a small fraction of existing cGLR diversity in animals. To discover new cGLR pattern recognition receptors in animal immunity, we coupled kingdom-wide bioinformatics analysis with a large-scale, forward biochemical screen of recombinant enzymes. Building on previous analysis of structurally and functionally related CD-NTase enzymes in bacteria that control anti-phage defense (Whiteley et al., 2019; Cohen et al., 2019), we constructed a hidden Markov model and used iterative PSI-BLAST to search all published metazoan genomes for proteins with homology to human cGAS, *Drosophila* cGLR1, or a structurally characterized cGLR from the beetle species *Tribolium castaneum* (Slavik et al., 2021). Resulting protein sequences were sorted into clusters using MMSeq2 and manually curated to identify high confidence cGLR enzymes according to three criteria: (1) conservation of the active site h[QT]GS [X8–20] [DE]h [DE]h [X50–90] h[DE]h motif known to be essential for nucleotide second messenger synthesis (Sun et al., 2013; Gao et al., 2013a; Civril et al., 2013; Kranzusch et al., 2014), (2) shared sequence homology across both the N-terminal NTase core and C-terminal helix-bundle required to complete the caged architecture of cGLR and CD-NTase enzymes (Kranzusch, 2019), and (3) predicted direct structural homology to known cGLR and CD-NTase enzymes based on AlphaFold2 modeling of representative protein sequences from each cluster (Jumper et al., 2021). The final curated list of animal cGLRs with putative active sites includes 3,020 unique enzymes conserved across 583 species (Figure 1A; Table S1). cGLRs are present in nearly all metazoan phyla with most metazoan genomes encoding 1–5 unique representatives (Figure 1A, B; Table S1). Animal species in some metazoan phyla, notably cnidarians and bivalves like *Stylophora pistillata* (stony coral) and *Crassostrea gigas* (Pacific oyster), encode a high number of cGLRs in their genomes. In some bivalves such as *Dreissena polymorpha* (Zebra mussel) and *Crassostrea virginica* (Eastern oyster), extreme radiation of *cGLR* genes results in >200 cGLR proteins encoded in a single genome (Figure 1A, B; Table S1). In the human genome, cGLRs with complete putative active sites include cGAS and MB21D2 (C3orf59), a cGLR previously identified by structural homology (Slavik et al., 2021) and known to be frequently mutated in cancer (Campbell et al., 2016), and two nearly identical proteins Mab21L1 and Mab21L2. Previous structural analysis of Mab21L1-related proteins confirms homology with cGLR enzymes in innate immunity (de Oliveira Mann et al., 2016), but Mab21L1-related proteins have a non-immune role in developmental tissue patterning and it is not understood if these proteins synthesize nucleotide second messenger signals (Chow et al., 1995; Yamada et al., 2003; Rainger et al., 2014; Horn et al., 2015). Notably, some specific metazoan phyla such as nematodes and most platyhelminthes encode no predicted cGLRs other than Mab21-like proteins, suggesting that innate immune cGLR signaling pathways have been specifically lost in select metazoan lineages similar to genetic loss events observed for TLRs in rotifera and platyhelminthes and NLRs in *Drosophila* and *C. elegans* (Zhang et al., 2010; Gerdol et al., 2017; Leulier and Lemaitre, 2008; Kangale et al., 2021).

**Figure 1.**
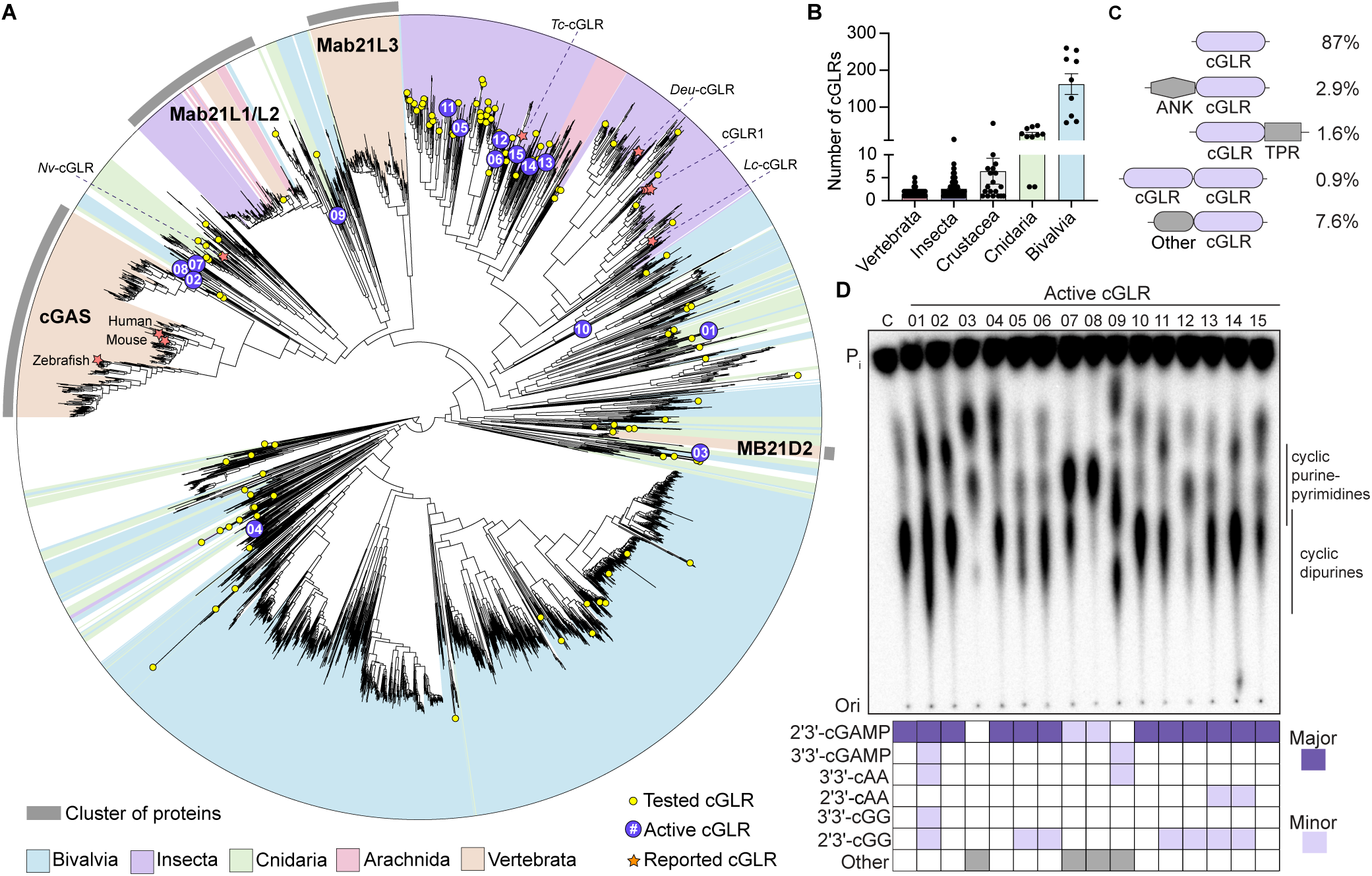
cGLRs are a widespread family of signaling enzymes in animal immunity. (A) Bioinformatic identification and phylogenetic tree of ∼3,000 predicted cGLRs from nearly all major animal phyla. Characterized cGLRs including vertebrate cGAS proteins, *N. vectensis* cGLR, and *Drosophila* cGLR1 are denoted with an orange star. Closely related *Mab21L1/2/3* genes are included but do not contain a predicted functional active site. Yellow circles represent cGLRs cloned for the biochemical screen, and purple circles denote active cGLRs selected for in-depth analysis. For additional information, see Figure S3 and Table S1. (B) Number of *cGLR* genes encoded in individual species categorized by animal phylum. (C) Domain organization of cGLR proteins. The prevalence of each domain architecture in sequenced animal genomes is listed as a percentage of all cGLR proteins (ANK = ankyrin repeat motif, TPR = tetratricopeptide repeat motif). Domain architectures that account <0.5% of all cGLR proteins are represented as “Other”. For additional details, see Table S2. (D) Thin layer chromatography and LC-MS/MS analysis of cGLR nucleotide second messenger products. cGLRs that synthesize a major product that could not be matched to a known nucleotide second messenger are denoted as “unknown”. Human cGAS was used as a control and was denoted as “C”; active cGLR number designations correspond to labels in Figure 1A. Data are representative of n = 3 independent experiments.

Analysis of cGLR sequences reveals specific patterns of evolution in animal immunity and protein features distinct from bacterial CD-NTase anti-phage defense enzymes. In most instances, cGLRs encoded within individual animal genomes map to disparate parts of the cGLR protein family tree demonstrating significant divergence and likely control of unique signaling pathways (Figure 1A and S1). An exception is insect genomes which often encode clusters of related cGLR enzymes that likely arose through recent gene amplification and diversification (Figure 1A and S1) (Slavik et al., 2021; Holleufer et al., 2021). In agreement with a role in activating innate immunity through synthesis of nucleotide second messenger signals, the vast majority of cGLRs (87%) are encoded as individual enzymatic proteins with no additional appended signaling domains (Figure 1C). However, some cGLRs contain predicted protein interaction domains including tetratricopeptide- and ankyrin-repeat modules also observed in TLR and RLR signaling pathways and other cGLRs are encoded as multi-cGLR domain fusions similar to OAS2 and OAS3 signaling enzymes in mammalian immunity (Figure 1C and S2A, Table S2) (Hur, 2019).

To investigate the function of metazoan cGLRs, we selected 140 representative sequences from across the cGLR phylogenetic tree and purified each recombinant enzyme for biochemical analysis (Figure 1A and S3, Table S3). These proteins are highly divergent in amino acid sequence and represent 72 species broadly distributed across the animal kingdom. For initial screening, each recombinant protein was incubated with a pool of common nucleic acid PAMPs as potential activating ligands, and nucleotide second messenger synthesis was analyzed using radiolabeled nucleotide substrates and thin-layer chromatography (TLC) (Figure S3). Consistent with a predicted role in innate immunity, many recombinant cGLRs robustly synthesized cyclic nucleotide products *in vitro* similar to human cGAS and *Drosophila* cGLR1 (Figure 1D and S3). We next selected 15 active cGLRs (Table S3) for in-depth analysis and used reactions containing individual combinations of NTPs to determine a nucleotide substrate-dependency profile for each enzyme (Figure S4). Combined with liquid chromatography-tandem mass spectrometry (LC-MS-MS) analysis, these results demonstrate that the most frequently detected cGLR product is the known nucleotide second messenger 2′3′-cGAMP, supporting a clear role for 2′3′-cGAMP as a common immune signaling molecule throughout the animal kingdom. 2′3′-cGAMP and 3′2′-cGAMP are the only cGLR-produced cyclic dinucleotide signals previously known to be synthesized in animals (Duncan-Lowey and Kranzusch, 2022). Intriguingly, nucleotide incorporation and LC-MS-MS analysis demonstrated that several cGLRs including cGLR-03, -07, -08, and -09 in our screen synthesize cyclic nucleotide products that do not match any known animal cGLR or bacterial CD-NTase standard, revealing that animal cGLRs are capable of synthesizing further diverse nucleotide second messengers in response to PAMPs (see below). Together these results reveal cGLRs as a widely distributed family of pattern recognition receptors in animals and define a model panel of active enzymes to establish the general rules of cGLR immune signaling.

### Divergent metazoan cGLRs respond to specific dsRNA and dsDNA nucleic acid PAMPs

To define the PAMP ligands sensed by animal cGLRs, we next determined the specific requirements for activation of each cGLR in our model panel of 15 enzymes. First, we confirmed via targeted mutagenesis of four representative enzymes that nucleotide second messenger synthesis is dependent on the putative cGLR active site motif (Figure S5A). The only known cGLR activating ligands are double-stranded DNA (human cGAS and close vertebrate homologs) (Sun et al., 2013) and double-stranded RNA (*Drosophila* cGLR1 and *T. castaneum* cGLR) (Slavik et al., 2021; Holleufer et al., 2021). We therefore next performed a series of reactions monitoring activity of each cGLR either alone, in the presence of a 45 bp immunostimulatory dsDNA (Stetson and Medzhitov, 2006), or the immunostimulatory dsRNA mimic polyI:C. Two cGLRs (cGLR-07 and -08) from the oyster species *C. gigas* and *C. virginica* were specifically activated by dsDNA, and seven cGLRs from diverse species were specifically activated by dsRNA (Figure 2A, B and S5B). Activation of each dsDNA/dsRNA-sensing cGLR was length-dependent, with longer nucleic acids being required for robust enzyme activation similar to previous results observed with human cGAS and *Drosophila* cGLR1 (Figure 2C and S5C, D) (Andreeva et al., 2017; Zhou et al., 2018; Slavik et al., 2021). The dsDNA-sensing cGLRs map closely to human cGAS within the cGLR protein family tree. In contrast, the dsRNA-sensing cGLRs are notably divergent and reside within distinct branches (Figure 1A and S1). Overall, these results demonstrate that recognition of foreign nucleic acid is a broadly shared mechanism of cGLR activation in metazoans and support that dsDNA-sensing may be a relatively recent evolutionary adaptation compared to dsRNA-recognition.

**Figure 2.**
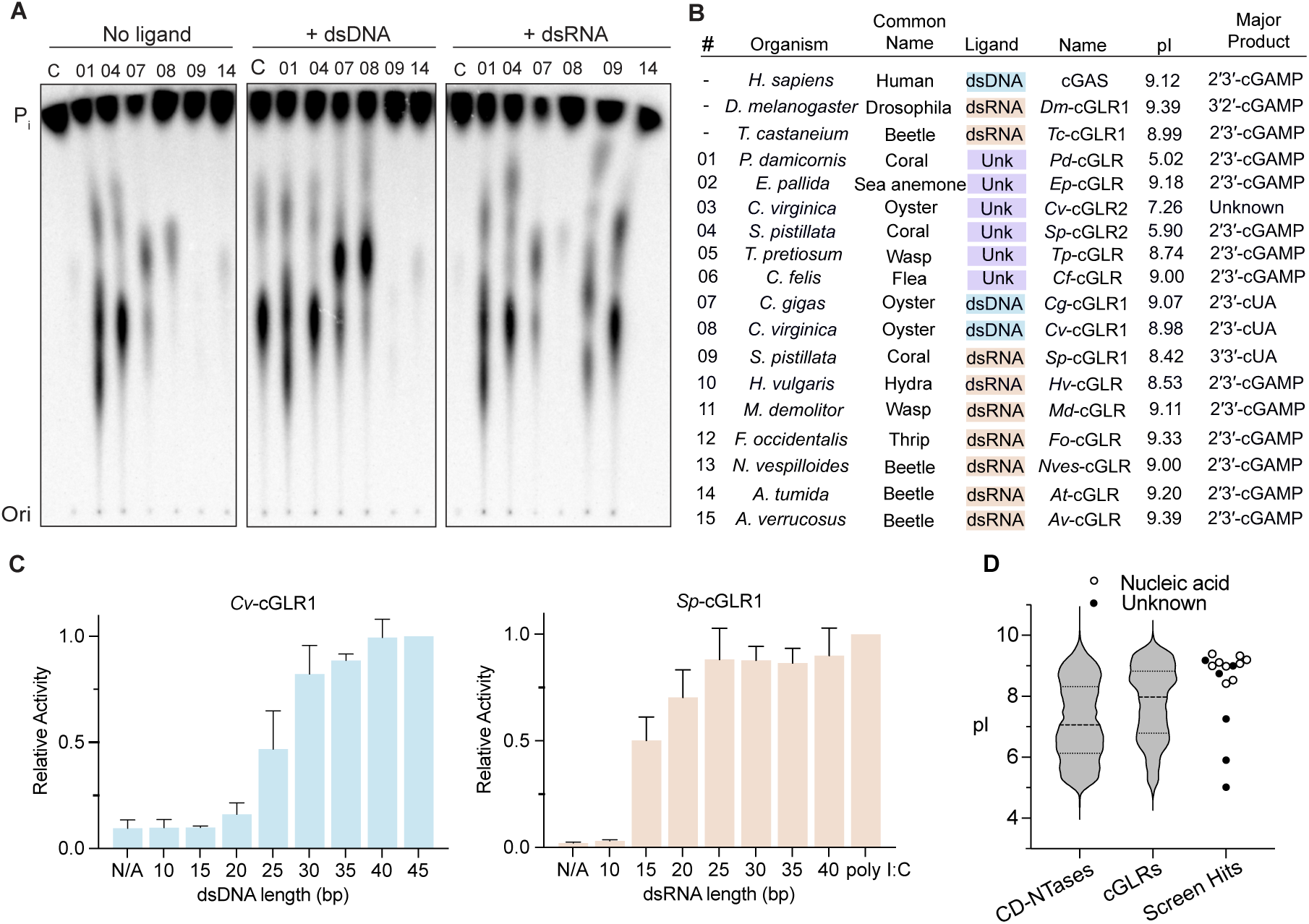
Divergent metazoan cGLRs respond to the common PAMPs dsDNA and dsRNA. (A) Thin layer chromatography analysis of animal cGLR activation in the presence of a 45 bp dsDNA or polyI:C dsRNA. Human cGAS was used as a control and was denoted as “C”; active cGLR number designations correspond to labels in Figure 1A. Data are representative of n = 3 independent experiments. (B) Summary of specific activating ligand and nucleotide second messenger product details for known cGLRs and each active cGLR enzyme identified here. See Figure 3 for analysis of nucleotide second messenger product identification for cGLR-07, -08, and -09. (C) *Cv*-cGLR1 and *Sp*-cGLR1 *in vitro* activity was analyzed in the presence of a panel of dsDNA or dsRNA activating ligands. Data are the mean ± std of n = 3 independent experiments. (D) Comparative analysis of predicted isoelectric point (pI) of bacterial CD-NTases, animal cGLRs, and new active animal cGLR enzymes demonstrates that nucleic acid sensing cGLRs (circles) are predicted to have a more positively charged surface compared to auto-active cGLRs (black dots) and bacterial CD-NTases. Full data on predicted pI of cGLRs and bacterial CD-NTases are included in Table S4.

An intriguing subset of cGLRs identified in our screen is a group of six enzymes that are robustly active in the absence of exogenously provided ligand. Consistent with recognition of negatively charged nucleic acid molecules, dsDNA- and dsRNA-sensing cGLRs exhibit a high isoelectric point (pI) (>8.5) and are predicted to have long, positively charged primary ligand binding surfaces similar to human cGAS and *Drosophila* cGLR1 (Figure 2D, S2B and S5E). The auto-active cGLR-02 from *E. pallida* (*Ep-*cGLR) fits this pattern and maps closely with cGAS and the oyster dsDNA-sensing cGLRs (Figure 1A and 2B), suggesting that in some cases observed auto-activity may be due to residual nucleic acid co-purified during protein purification. In contrast, some auto-active cGLR enzymes are highly acidic with cGLR-01 from *P. damicornis* (*Pd-*cGLR) for example exhibiting a calculated isoelectric point as low as 5.0 (Figure 2B). Structural modeling of acidic cGLRs predicts a negatively charged primary ligand-binding groove consistent with the hypothesis that these enzymes respond to a positively charged ligand, not nucleic acid, that may have co-purified during recombinant protein purification from bacteria (Figure S5E). These data support that cGLRs are likely capable of responding to yet unknown non-nucleic acid ligands, similar to TLRs recognizing structurally divergent PAMPs including nucleic acid, LPS, and bacterial flagellin protein (Kawai and Akira, 2010).

### Metazoan cGLRs produce diverse cyclic di-purine and purine-pyrimidine signals

Following ligand binding and enzyme activation, cGLRs catalyze synthesis of a nucleotide second messenger product to control downstream immune signaling (Sun et al., 2013; Slavik et al., 2021). In addition to vertebrate cGAS enzymes, 2′3′-cGAMP has previously been identified as the nucleotide product of a cGLR from the beetle insect *T. castaneum* (Slavik et al., 2021), and the cnidarian *Nematostella vectensis* (Kranzusch et al., 2015; Gui et al., 2019). Analysis of our screen of animal cGLR enzymes extends 2′3′-cGAMP production to a range of species in diverse major animal phyla including corals (*P. damicornis*), hydras (*H. vulgaris*), fleas (*C. felis*), and thrips (*F. occidentalis*) (Figure 2B).

Several active cGLR enzymes in our screen including cGLRs -07, -08, and -09 (*Cg*-cGLR1, *Cv*-cGLR1 and *Sp*-cGLR1) produce nucleotide second messenger products that exhibit a distinct migration pattern on TLC compared to 2′3′-cGAMP (Figure 1D). To identify the nucleotide products of these enzymes, we incubated *Cg*-cGLR1, *Cv*-cGLR1 and *Sp*-cGLR1 reactions with each individual α^32^P-radiolabeled NTPs (ATP, GTP, CTP, UTP) and combinations of non-radiolabeled NTPs (Figure 3A and S4). The major products of *Cg*-cGLR1, *Cv*-cGLR1, and *Sp*-cGLR1 were each specifically labeled with adenosine and uridine, and were resistant to phosphatase treatment that removes terminal phosphate groups, suggesting synthesis of hybrid purine-pyrimidine cyclic dinucleotide signals (Figure 3A). Using nucleobase-specific labeling and specific digestion of 3′−5′ phosphodiester bonds with Nuclease P1, we observed that the nucleotide products of *Cg*-cGLR1 and *Cv*-cGLR1 contain a 3′−5′ linkage incorporating the adenosine phosphate and a protected 2′−5′ linkage incorporating the uridine phosphate, indicating synthesis of a mixed-linkage 2′–5′ / 3′–5′ cyclic UMP–AMP species (2′3′-cUA) (Figure 3A). In contrast, we observed that the nucleotide product of *Sp*-cGLR1 contains only 3′−5′ linkages indicating synthesis of a canonically linked product 3′–5′ / 3′–5′ cyclic UMP–AMP (3′3′-cUA) (Figure 3A). To verify these findings, we chemically synthesized 2′3′-cUA and 3′3′-cUA as synthetic standards and used comparative high-performance liquid chromatography (HPLC), tandem mass spectrometry profiling (MS/MS), and nuclear magnetic resonance spectroscopy (NMR) to confirm that the dsDNA-activated *Cg*-cGLR1 and *Cv*-cGLR1 enzymes produce the unique nucleotide signal 2′3′-cUA, and the dsRNA-activated *Sp*-cGLR1 enzyme produces the nucleotide signal 3′3′-cUA (Figure 3B, C and S6). These results demonstrate that animal cGLRs can incorporate pyrimidine bases to diversify innate immune signaling products and reveal the first metazoan enzymes capable of synthesizing purine-pyrimidine hybrid cyclic dinucleotides.

**Figure 3.**
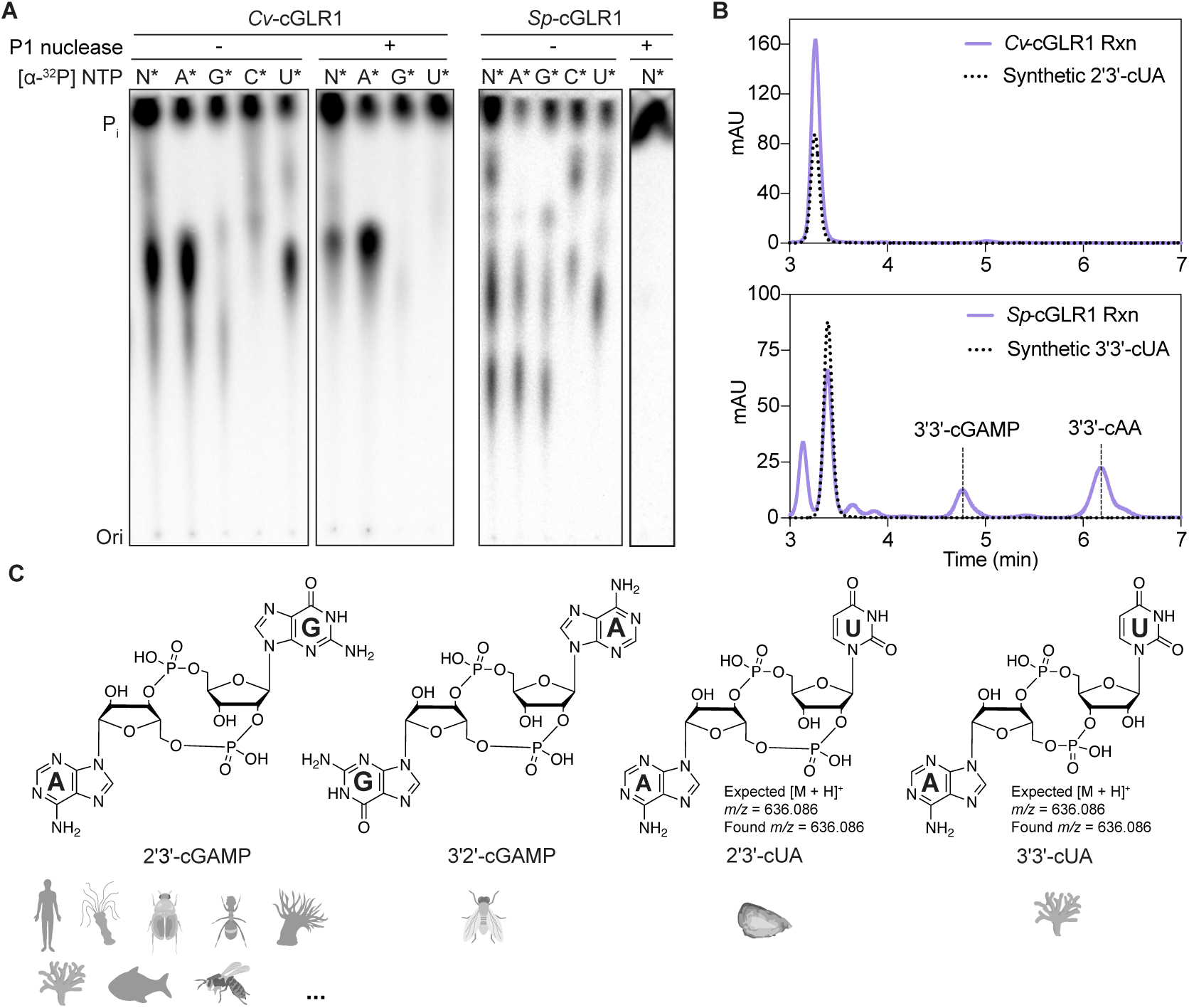
Metazoan cGLRs produce diverse cyclic di-purine and purine-pyrimidine signals. (A) Thin layer chromatography analysis of *Cv*-cGLR1 and *Sp*-cGLR1 reactions labeled with individual α^32^P-NTPs and treated as indicated with nuclease P1 (a 3′–5′ phosphodiester bond-specific nuclease) and CIP (a phosphatase that removes terminal phosphate groups from nucleotides). Nucleotide labeling and P1 digestion suggest that *Cv*-cGLR1 and *Sp*-cGLR1 synthesize 2′3′-cUA and 3′3′-cUA. Data are representative of n = 3 independent experiments. (B) HPLC analysis *Cv*-cGLR1 and *Sp*-cGLR1 reactions compared to 2′3′-cUA and 3′3′-cUA synthetic standards. Data are representative of n = 3 independent experiments. (C) LC-MS/MS and NMR verification of the *Cv*-cGLR1 and *Sp*-cGLR1 products as 2′3′-cUA and 3′3′-cUA. cGLR analysis supports that although 2′3′-cGAMP is the most common nucleotide second messenger product, cGLRs are capable of synthesizing diverse immune signals. See also Figure S6.

### Animals encode STING receptors with distinct nucleotide second messenger preferences

cGLR nucleotide second messenger signals are recognized in animal cells by the downstream receptor STING (Burdette et al., 2011; Sun et al., 2013; Kranzusch et al., 2015); (Slavik et al., 2021; Holleufer et al., 2021). To define how cGLRs control immune activation, we next extended our bioinformatic analysis to map STING receptor diversity in metazoans (Figure 4A,B, Table S5). Humans, as well as the model animals zebrafish and *Drosophila*, encode one copy of STING (Figure 4A). Moreover, we found that in most cases (88%) animal genomes encode only one STING receptor, suggesting that STING often functions as a shared signaling adapter for multiple endogenous cGLR signaling pathways (Figure 4B, Table S5). However, we found evidence for more complex signaling networks in some animal species such as the coral *S. pistillata* and the oyster *C. virginica*, where 3–10 independent *STING* genes are encoded in genomes exhibiting extreme radiation of >20 *cGLR* genes (Figure 4B).

**Figure 4.**
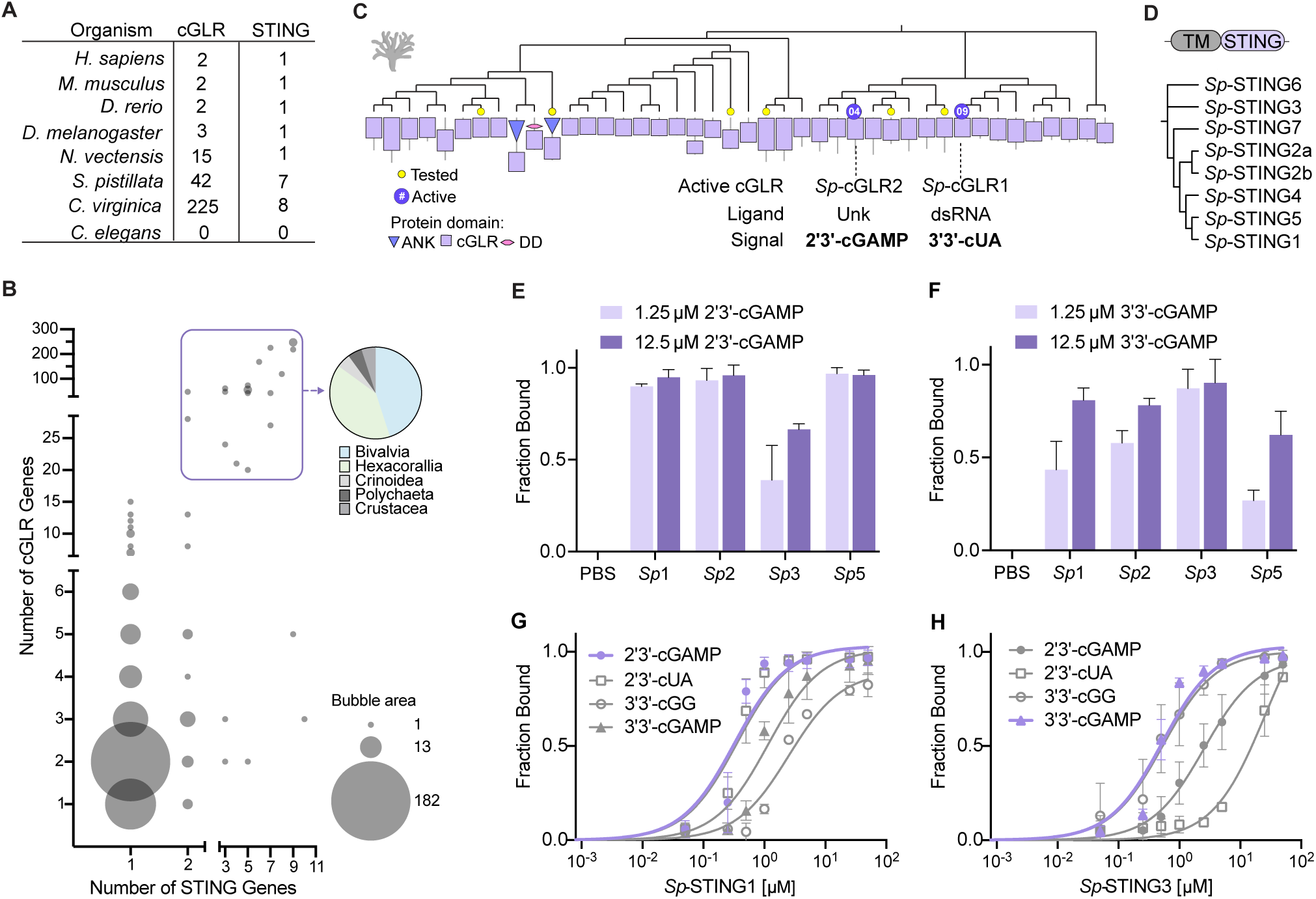
Animals encode STING receptors with distinct nucleotide second messenger preferences. (A) cGLR and STING protein diversity in select representative organisms from different animal phyla. Mab21L1-like proteins are excluded. (B) Analysis of *cGLR* and *STING* gene copy number in 381 animal species. Bubble size corresponds to species frequency. Animal genomes demonstrating significant *cGLR* and *STING* gene radiation are highlighted in a purple box and the derived taxa are plotted in the corresponding pie chart. Full data of the number of cGLR and STING genes in each animal species are included in Table S5. (C) Diversity of cGLR proteins in the stony coral *S. pistillata* including two active enzymes *Sp*-cGLR1 and *Sp*-cGLR2 identified in the biochemical screen. *S. pistillata* cGLRs fused to ankyrin repeats (ANK) and death domains (DD) potentially involved in protein–protein and protein–ligand interactions are annotated. *Sp*-cGLR1 produces 3′3′-cUA in response to dsRNA and *Sp*-cGLR2 produces 2′3′-cGAMP in response to an unknown ligand. (D) *S. pistillata* encodes seven STING proteins, each predicted to contain an N-terminal transmembrane domains (TM) and a C-terminal cyclic dinucleotide binding domain (CBD). *Sp*-STING proteins share ∼54–75% CBD sequence identity. (E,F) Quantification of electrophoretic mobility shift assay (EMSA) analysis of the binding of four *Sp*-STING receptors to 3′3′-cGAMP and 2′3′-cGAMP. Data are the mean ± std of n = 2 independent experiments. See also Figure S7A–C. (G,H) Quantification of EMSA analysis of affinity of *Sp*-STING1 and *Sp*-STING3 with the nucleotide second messengers 2′3′-cGAMP, 2′3′-cUA, 3′3′-cGG and 3′3′-cGAMP demonstrates that *Sp*-STING1 preferentially binds 2′3′-linked cyclic dinucleotides and *Sp*-STING3 preferentially binds 3′3′-linked cyclic dinucleotides. Results are plotted as fraction bound (shifted/total signal) as a function of increasing protein concentration and fit to a single binding isotherm. Data are the mean ± std of n = 2 or 3 independent experiments. See also Figure S7D.

To understand how animal cGLRs may control more complex signaling networks, we focused on biochemical analysis of *S. pistillata* cGLR-STING signaling pathways. *S. pistillata* encodes 42 cGLRs and 7 STING receptors (Figure 4C,D). *S. pistillata* cGLRs are highly variable with some pairs of enzymes sharing <25% sequence identity and 3 cGLR proteins with the core NTase domain fused to additional ligand-interacting domains including ankyrin repeats and death domains (Figure 4C). Two active *S. pistillata* cGLRs were identified in our biochemical screen, *Sp*-cGLR1 a dsRNA-sensing enzyme that produces the nucleotide signal 3′3′-cUA and *Sp*-cGLR2 a 2′3′-cGAMP synthesizing enzyme activated by an unknown ligand, demonstrating that one organism can encode cGLRs synthesizing distinct 3′3′-or 2′3′-linked nucleotide second messenger signals (Figures 2 and 3). The CDN-binding domain of *S. pistillata* STING receptors are 54–75% identical at the amino acid level, and each protein has an architecture similar to human and *Drosophila* STING with an N-terminal transmembrane domain fused to a C-terminal cyclic dinucleotide binding domain (Figure 4D). We successfully purified the STING cyclic dinucleotide binding domain from four *S. pistillata* STING proteins (*Sp*-STING1, *Sp*-STING2, *Sp*-STING3, and *Sp*-STING5) and used an electrophoretic mobility shift assay to measure the ability of each of these receptors to bind 3′3′- or 2′3′-linked cyclic dinucleotide signals. Each *S. pistillata* STING protein exhibited a unique pattern of cyclic dinucleotide affinity, suggesting that cGLR-dependent synthesis of specific nucleotide second messengers can lead to distinct profiles of receptor activation (Figure 4E–F and S7A). Three *Sp*-STING proteins preferentially bound 2′3′-linked cyclic dinucleotides including 2′3′-cGAMP and 2′3′-cUA, consistent with the typical preference of metazoan STING receptors to bind noncanonically-linked cyclic dinucleotide signals (Figure 4E–F, S7B) (Ablasser et al., 2013; Kranzusch et al., 2015). In contrast, *Sp*-STING3 demonstrated clear preference for 3′3′-linked cyclic dinucleotides including the bacterial nucleotide second messengers 3′3′-cGAMP and 3′3′-cGG (Figure 4E–F, S7B). Together these results demonstrate that animal STING receptors can exhibit specific nucleotide second messenger preferences and suggest that expression of multiple STING proteins creates discrete cGLR-STING signaling networks in animal cells.

### Molecular mechanism of STING ligand recognition in *S. pistillata*

To define the molecular mechanism of preferential recognition of 3′3′- and 2′3′-linked nucleotide second messengers in *S. pistillata* cGLR-STING signaling, we determined crystal structures of the *Sp*-STING3–3′3′-cGAMP complex (1.7 Å) and the *Sp*-STING1–2′3′-cGAMP complex (2.1 Å) (Figure 5A and Table S6). Similar to human STING and previous structures of metazoan STING proteins (Gao et al., 2013b; Zhang et al., 2013; Kranzusch et al., 2015; Morehouse et al., 2020; Slavik et al., 2021), both *Sp*-STING proteins form a characteristic V-shaped, homodimeric receptor domain architecture that binds cyclic dinucleotide ligands within a deep central pocket (Figure 5A). Previous structures of STING–cyclic dinucleotide complexes demonstrate that high-affinity recognition of endogenous cGLR signals results in closure of the STING β-strand lid domain and rotation at the dimer interface to create a tightly compact configuration with ∼32–36 Å distance between the top of each STING protomer (Zhang et al., 2013; Slavik et al., 2021). In contrast, STING binding to lower affinity cyclic dinucleotide ligands does not induce subunit rotation and results in a more open structural state with an ∼47–55 Å distance between protomers that is associated with weaker downstream signaling (Gao et al., 2013b; Zhang et al., 2013; Kranzusch et al., 2015; Ergun et al., 2019). Both *Sp*-STING3–3′3′-cGAMP and *Sp*-STING1–2′3′-cGAMP complexes exhibit a fully compact state with the β-strand lid domain closed and rotation along the dimer interface resulting in 30.9 and 30.6 Å between the top of each STING protomer (Figure 5A). The compact *Sp*-STING3–3′3′-cGAMP and *Sp*-STING1–2′3′-cGAMP structural states are superimposable with the human STING–2′3′-cGAMP and *Drosophila* STING–3′2′-cGAMP complexes supporting that *Sp*-STING proteins recognize 3′3′- and 2′3′-linked cyclic dinucleotides as high-affinity, endogenous cGLR immune signals. Receptor domain rotation and structural compaction upon cyclic dinucleotide recognition is an evolutionarily ancient feature of STING signaling, with bacterial STING receptors in anti-phage defense systems adopting a similar closed compact configuration when bound to the 3′3′-linked linked CD-NTase signal 3′3′-c-di-GMP (Morehouse et al., 2020; Morehouse et al., 2022). Notably, *Sp*-STING3 is the first example of a metazoan STING receptor capable of preferentially recognizing canonically-linked cyclic dinucleotides with high affinity and adopting a compact, fully active state in complex with a 3′3′-linked nucleotide second messenger (Figure 5A).

**Figure 5.**
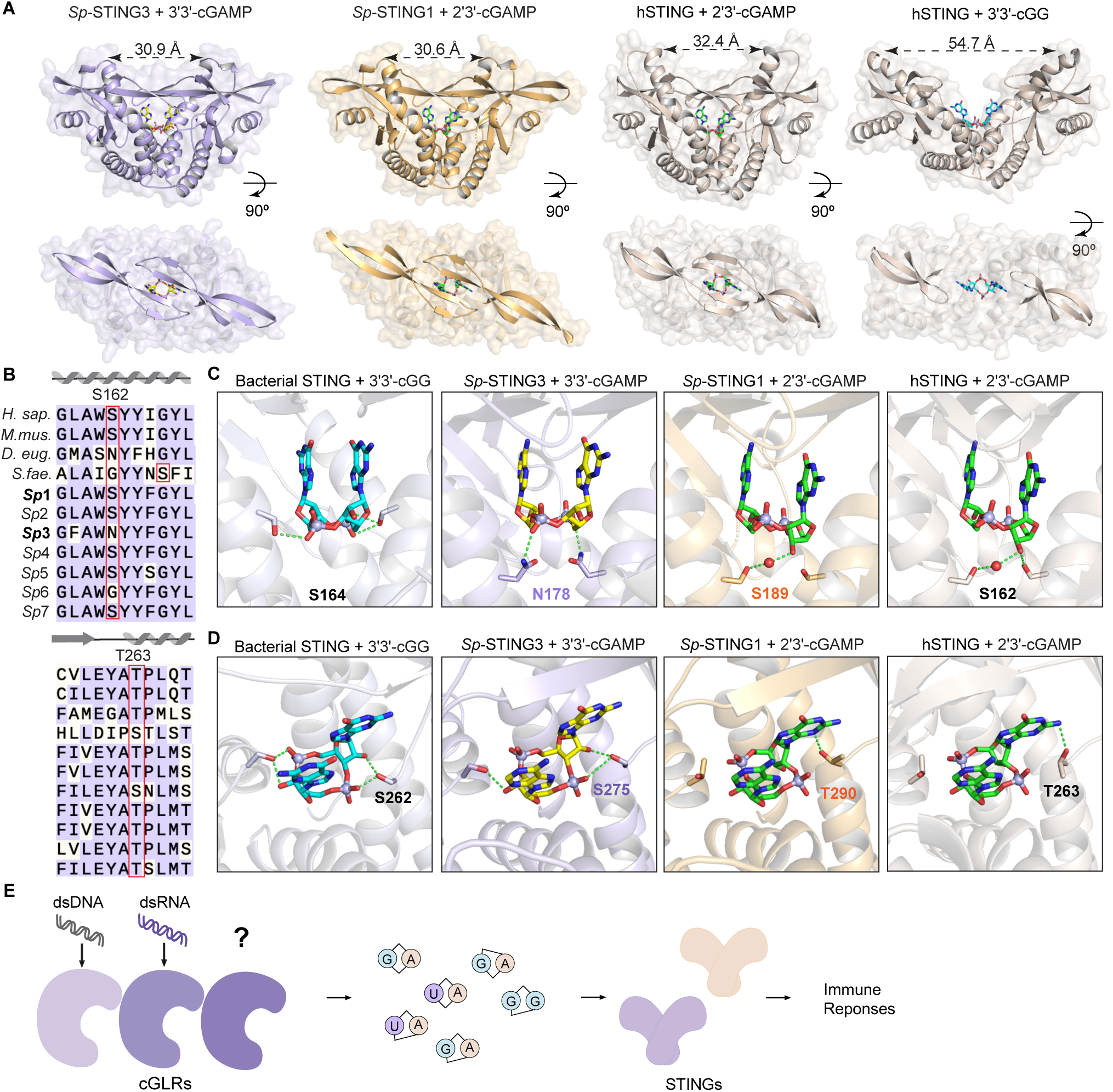
Molecular mechanism of STING ligand recognition in *S. pistillata*. (A) Crystal structures of *Sp*-STING1 in complex with 2′3′-cGAMP and *Sp*-STING3 in complex with 3′3′-cGAMP. The *Sp*-STING1–2′3′-cGAMP and *Sp*-STING3–3′3′-cGAMP structures adopt a tightly closed conformation with an ordered β-strand lid most similar to the human STING–2′3′-cGAMP complex, supporting high-affinity recognition of an endogenous cGLR nucleotide second messenger signal. (B) Sequence alignment of *Sp*-STING receptors with STING proteins from representative animal species and the bacterium *S. faecium* reveals that *Sp*-STING3 has unique substitutions at key residues involved in cyclic dinucleotide binding. (C,D) Structural comparison of the bacterial *Sf*-STING–3′3′-cGG complex, *Sp*-STING3–3′3′-cGAMP complex, *Sp*-STING1–2′3′-cGAMP complex and human STING–2′3′-cGAMP complex reveals critical difference in phosphodiester linker recognition that explain specificity for 2′3′-linked and 3′3′-linked cGLR nucleotide second messenger signals. (E) Model of cGLR signaling pathway in animal immunity. Animals encode multiple cGLRs that recognize diverse PAMPs and produce distinct nucleotide second messengers. STING receptor duplication and cyclic dinucleotide-specific adaptations enables creation of complex cGLR-STING signaling networks.

We next analyzed the ligand binding pockets of *Sp*-STING receptors compared to the human STING–2′3′-cGAMP and bacterial *S. faecium* STING–3′3′-c-di-GMP complex structures. *Sp*-STING3 and *Sp*-STING1 encode a pair of conserved aromatic residues (*Sp*-STING3 Y183 and Y253, *Sp*-STING1 Y194 and Y263) that stack against each nucleobase of the ligand (Figure S8A,B). Additionally, closure of the β-strand lid domain positions *Sp*-STING3 R250 and *Sp*-STING1 R261 to make direct contacts with the ligand phosphate backbone or nucleobase face similar to recognition of cyclic dinucleotides by human STING R238 and bacterial *S. faecium* STING R234 (Figure S8A,B). Each of these major interactions are shared across all known bacterial and metazoan STING receptors and define universally conserved aspects of STING cyclic dinucleotide recognition (Morehouse et al., 2020). Although nucleobase stacking and β-strand lid domain interactions are highly conserved, key contacts at the bottom of the STING ligand-binding pocket diverge between *Sp*-STING3 and *Sp*-STING1 and explain altered ligand specificity. First, in the *Sp*-STING1–2′3′-cGAMP structure, residue S189 interacts with the guanosine 3′-OH to coordinate the phosphate and ribose ring similar to interactions between human STING S162 and 2′3′-cGAMP (Figure 5B,C) (Zhang et al., 2013). In contrast, in *Sp*-STING3 an asparagine substitution N178 at the equivalent position extends the side chain further into the binding pocket and allows direct interactions with the 3′3′-cGAMP phosphate group (Figure 5C). Second, *Sp*-STING3 S275 on either side of the pocket interacts with the free 2′-OH of each nucleobase allowing specific coordination of the 3′3′-linked backbone of 3′3′-cGAMP (Figure 5B,D). In *Sp*-STING1 and human STING, threonine residues at this position are unable to access a similar rotamer conformation and instead contact the guanosine base of 2′3′-cGAMP (Figure 5B,D). Notably, each of the *Sp*-STING3–3′3′-cGAMP interactions specific to recognition of 3′3′-linked signals are mechanistically similar to how bacterial *S. faecium* STING coordinates the 3′3′-linked backbone of 3′3′-cGG (Morehouse et al., 2020; Morehouse et al., 2022). *S. faecium* STING S164 makes a phosphate-specific contact similar to *Sp*-STING3 N178, and *S. faecium* STING S262 is positioned with nearly identical 3′3′-linked backbone contacts as *Sp*-STING3 S276 (Figure 5B–D). Additionally, similar serine-to-asparagine substitutions in *Drosophila* STING are also responsible for a switch in ligand specificity from 2′3′-cGAMP to 3′2′-cGAMP (Slavik et al., 2021), further supporting that key serine and asparagine positions at the base of the STING ligand binding pocket control specificity for individual cGLR nucleotide second messenger signals. Together, these results explain the molecular mechanism of cGLR-STING signaling pathways in *S. pistillata* and provide a model for understanding evolution of cGLR nucleotide second messenger signal specificity.

## Discussion

Our data demonstrate that cGLRs are a diverse, widespread family of pattern recognition receptors conserved across nearly every metazoan phylum. Building upon the seminal discovery of cGAS as a dsDNA sensor in vertebrates (Sun et al., 2013) and the recent identification of cGLR1 as a dsRNA sensor in *Drosophila* (Slavik et al., 2021; Holleufer et al., 2021), our forward biochemical screen reveals thousands of diverse cGLRs conserved throughout the animal kingdom. We describe 15 new active cGLR enzymes with representatives broadly distributed in phylogenetically distant animals including insects, anemones, mollusks, hydras, and corals that respond to common long double-stranded nucleic acid PAMPs. Similar to other major families of pattern recognition receptors including TLRs, RLRs, NLRs, and CLRs (Pandey et al., 2014; Tan et al., 2018; Wang et al., 2021; Geijtenbeek and Gringhuis, 2009), cGLRs likely form a network of signaling pathways that enable animal cells to sense diverse microbial pathogens.

Large-scale biochemical analysis of cGLRs defines rules that control this type of signaling pathway in animal innate immunity (Figure 5E). Animal cGLRs typically reside in an inactive state and require recognition of foreign ligands to initiate signaling. We show that many diverse metazoan cGLRs respond to the PAMPs dsDNA and dsRNA (Figure 2), demonstrating that a common initial step in cGLR activation is sensing of nucleic acid molecules associated with pathogen replication. Biochemical analysis and structural modeling support previous findings that cGLRs share a conserved overall protein architecture where PAMP recognition occurs within a primary ligand-binding surface on the back-face of the enzymatic nucleotidyltransferase domain (Slavik et al., 2021). Diversification of this primary cGLR ligand-binding surface, as typified by enzymes like *Pd*-cGLR and *Sp-*cGLR2 with ligand-binding surfaces predicted to be highly negatively charged (Figure S5E), likely enables animal cells to evolve cGLR signaling pathways with new specificities. Notably, some cGLRs appear auto-active in our biochemical screen and we hypothesize that this could be a result of activating bacterial ligands from *E. coli* introduced during protein purification. Following ligand stimulation, cGLRs become enzymatically active and catalyze synthesis of a nucleotide second messenger signal. In addition to synthesis of 2′3′-cGAMP as a common immune signal shared across many animals (Figures 1 and 3), we demonstrate that cGLRs are capable of synthesizing chemically diverse nucleotide second messengers using both purine and pyrimidine bases. *Cv*-cGLR1 and *Sp*-cGLR1 synthesis of the hybrid purine-pyrimidine cyclic dinucleotide signals 2′3′-cUA and 3′3′-cUA highlights how variable nucleobase- and phosphodiester linkage-specificity creates a large array of potential cGLR signaling products (Figure 3). Several previous studies have shown evidence of cGAS- or STING-related innate immune responses using *in vivo* or cell-based assays in individual invertebrate species (Kranzusch et al., 2015; Qiao et al., 2021; Li et al., 2022; Amparyup et al., 2021; Margolis et al., 2021). Our biochemical analysis now establishes a framework to define the molecular basis of activation and nucleotide second messenger signaling in these cGLR pathways.

Most animals encode 2–4 cGLR proteins and a single copy of the cyclic dinucleotide receptor STING. However, in some cases, like the coral *S. pistillata,* we observe dramatic expansion of these signaling pathways with >40 *cGLR* genes and 7 distinct STING receptors (Figure 4). Structures of *S. pistillata* STING proteins explain how alterations to the STING cyclic dinucleotide binding pocket can combine with the ability of cGLRs to synthesize diverse nucleotide signals to create a complex cGLR-STING signaling network within a single species (Figure 5). In vertebrates, activation of the cGAS-STING pathway leads to type-I IFN signaling, NF-κB-dependent transcription activation, and induction of autophagy responses that limit pathogen replication and tumorigenesis (Ablasser and Chen, 2019). Correspondingly, recent studies in insect, anemone, and choanoflagellate models suggest that cGLR-STING signaling in invertebrates triggers antiviral and anti-microbial resistance through NF-κB- and autophagy-dependent responses (Martin et al., 2018; Liu et al., 2018; Cai et al., 2020; Slavik et al., 2021; Holleufer et al., 2021; Margolis et al., 2021; Woznica et al., 2021; Gui et al., 2019). Animals encode additional receptors including RECON that have been identified as sensors for nucleotide second messenger signals (McFarland et al., 2017; Whiteley et al., 2019), suggesting that diverse cGLRs may also control STING-independent cellular responses.

The evolutionary connection between animal innate immunity and prokaryotic anti-phage defense provides an additional opportunity to expand understanding of conserved principles of cGLR biology in pathogen sensing and immune signaling. Previous structural and biochemical analysis of CD-NTase anti-phage defense enzymes in bacteria revealed diversification of nucleotide specificity, synthesis of pyrimidine-purine hybrid cyclic dinucleotides, and selective activation of immune receptors (Whiteley et al., 2019; Lowey et al., 2020)–all aspects that we now demonstrate are conserved features of animal cGLR signaling pathways. Most interestingly, human and other vertebrates encode uncharacterized cGLRs in addition to cGAS that have complete catalytic sites and likely control novel innate immune responses in cancer and other disease (Slavik et al., 2021). Large-scale biochemical discovery of cGLRs as a diverse family of PRRs provides a new foundation to expedite the understanding of these innate immune sensing pathways in mammalian immunity.

### Limitations of the Study

Our biochemical approach enabled discovery of diverse metazoan cGLR signaling enzymes that respond to known cGLR PAMPs including dsDNA and dsRNA nucleic acid. However, a key limitation of our screen is that many animal cGLR proteins appear inactive *in vitro* likely due to the requirement of these enzymes to be activated by recognizing yet unknown PAMPs. Previous analysis of the orphan cGLR MB21D2 in human cells demonstrates that this enzyme does not respond to a wide array of known PAMPs including the innate immune agonists LPS, bacterial lipopeptide, and yeast cell wall zymosan (Slavik et al., 2021). Future research will be required to define new ligands recognized by human MB21D2 and other animal cGLRs and explain why some cGLRs appear auto-active when purified from *E. coli*. Additionally, recognition of pathogen-associated nucleic acid molecules like dsRNA and dsDNA strongly suggests that newly identified cGLRs function in innate immunity similar to human cGAS and *Drosophila* cGLR1 (Sun et al., 2013; Slavik et al., 2021; Holleufer et al., 2021). Animal experiments will be essential to define these signaling pathways *in vivo* and fully explain the role of divergent cGLRs in the immune responses of non-model organisms.

## Acknowledgements

The authors are grateful to J. Eaglesham and members of the Kranzusch lab for helpful comments and discussion. The authors thank C. Deutscher for expert technical assistance with synthetic cyclic dinucleotide purifications and Dr. Yuan Fang (Boston University) for technical support with coloring scattered dot plot by density. The work was funded by grants to P.J.K. from the Pew Biomedical Scholars program, the Burroughs Wellcome Fund PATH program, the Richard and Susan Smith Family Foundation, The Mathers Foundation, The Mark Foundation for Cancer Research, the Parker Institute for Cancer Immunotherapy, the DFCI-Novartis Drug Discovery Program, and the National Institutes of Health (1DP2GM146250-01). Y.L. is supported as a Benacerraf Fellow in Immunology, K.M.S. is supported as an NCI F99 Graduate Fellow NIH 1F99CA274660-01, B.R.M. was supported as a Ruth L. Kirschstein NRSA Postdoctoral Fellow NIH F32GM133063, and C.C.d.O.M. was supported as a Cancer Research Institute/Eugene V. Weissman Fellow. X-ray data were collected at the Northeastern Collaborative Access Team beamlines 24-ID-C and 24-ID-E (P30 GM124165), and used a Pilatus detector (S10RR029205), an Eiger detector (S10OD021527) and the Argonne National Laboratory Advanced Photon Source (DE-AC02-06CH11357).

## Author Contributions

Experiments were designed and conceived by Y.L., K.M.S., B.R.M., C.C.d.O.M., and P.J.K. cGLR bioinformatic analysis and initial screen was performed by Y.L., K.M.S, B.R.M., C.C.d.O.M., and P.J.K. Biochemical and nucleotide second messenger analysis experiments were performed by Y.L. with assistance from K.M.S., B.R.M., and J.L. Crystallography experiments were performed by Y.L. with assistance from B.R.M. and K.M. Synthetic nucleotide product synthesis and characterization experiments were performed by D.K. and F.S. The manuscript was written by Y.L. and P.J.K. All authors contributed to editing the manuscript and support the conclusions.

## Declaration of Interests

D.K. and F.S. are employed at Biolog Life Science Institute GmbH & Co. KG, which sells 3′3′-cUA and may sell 2′3′-cUA as research tools.

## Figure Legends

## SI Figure Legends

**Figure S1.**
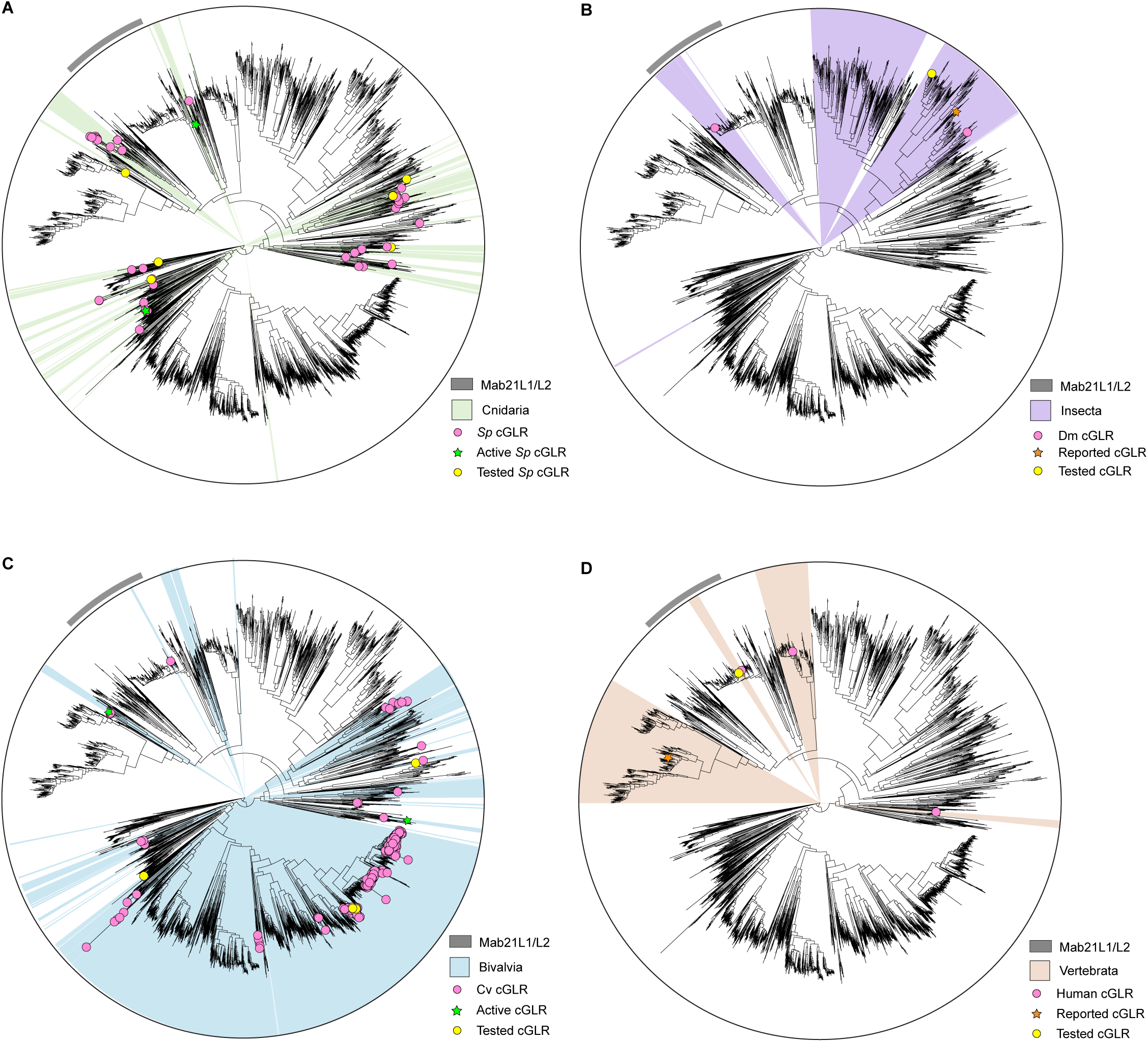
Divergence of cGLR in representative animal species, related to Figure 1. Analysis of cGLR diversity in (A) the cnidarian *S. pistillata*, (B) insect *D. melanogaster*, (C) bivalve *C. virginica*, and (D) human genome. Individual species encode diverse cGLR proteins from distinct parts of the protein family tree suggesting existence of distinct immune signaling pathways. One notable exception are insect genomes that typically encode clusters of closely related *cGLR* genes. Predicted cGLRs from bioinformatic analysis (pink), cGLRs tested in the biochemical screen (yellow) are denoted with a circle symbol. Active cGLRs identified in the biochemical screen (green) and previously reported active cGLRs (orange) are denoted with a star symbol.

**Figure S2.**
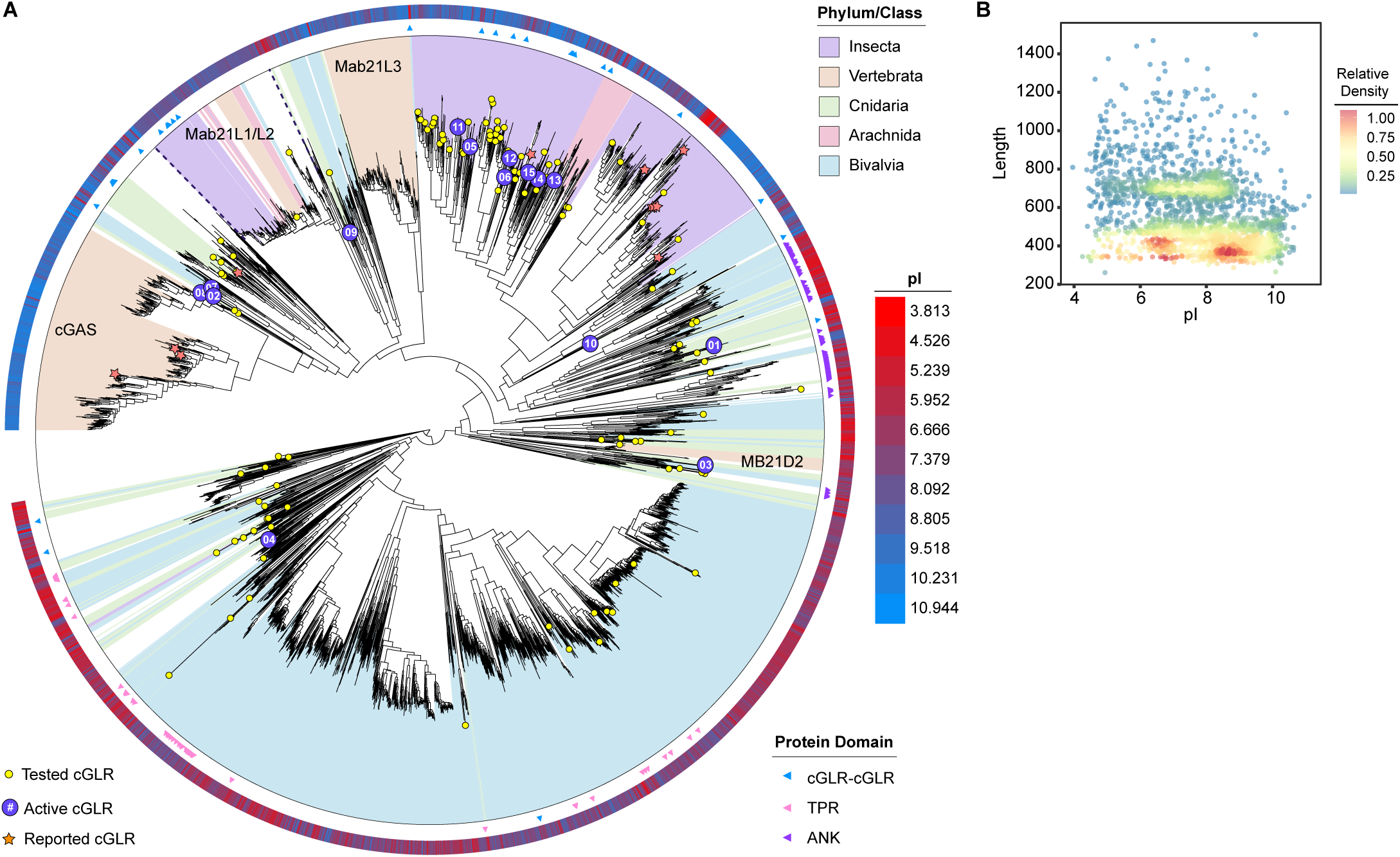
Analysis of cGLR protein domain architecture and predicted isoelectric point, related to Figure 1. (A) Analysis of cGLR protein domain architecture. Most cGLR proteins exist as single domains, but some are encoded as fusions to tetratricopeptide repeat (TPR) domains (pink), ankyrin-repeat (ANK) domains (purple), or as tandem connected cGLR domains (blue) and are denoted with a triangle symbol. The predicted isoelectric point for each cGLR protein is displayed as an outer colored ring ranging from negatively charged (red) to positively charged (blue). cGLR proteins that respond to dsDNA and dsRNA have a higher predicted isoelectric point supporting positive surface charge for interaction with negatively charged nucleic acid. (B) Calculated isoelectric point of cGLRs plotted against length of proteins reveals the presence of three major types, suggesting divergence of cGLRs in PAMP recognition. Color scale reflects density of dots, where red indicates high density and blue represents low density. Full data of protein length and calculated isoelectric points of cGLRs are included in Table S4.

**Figure S3.**
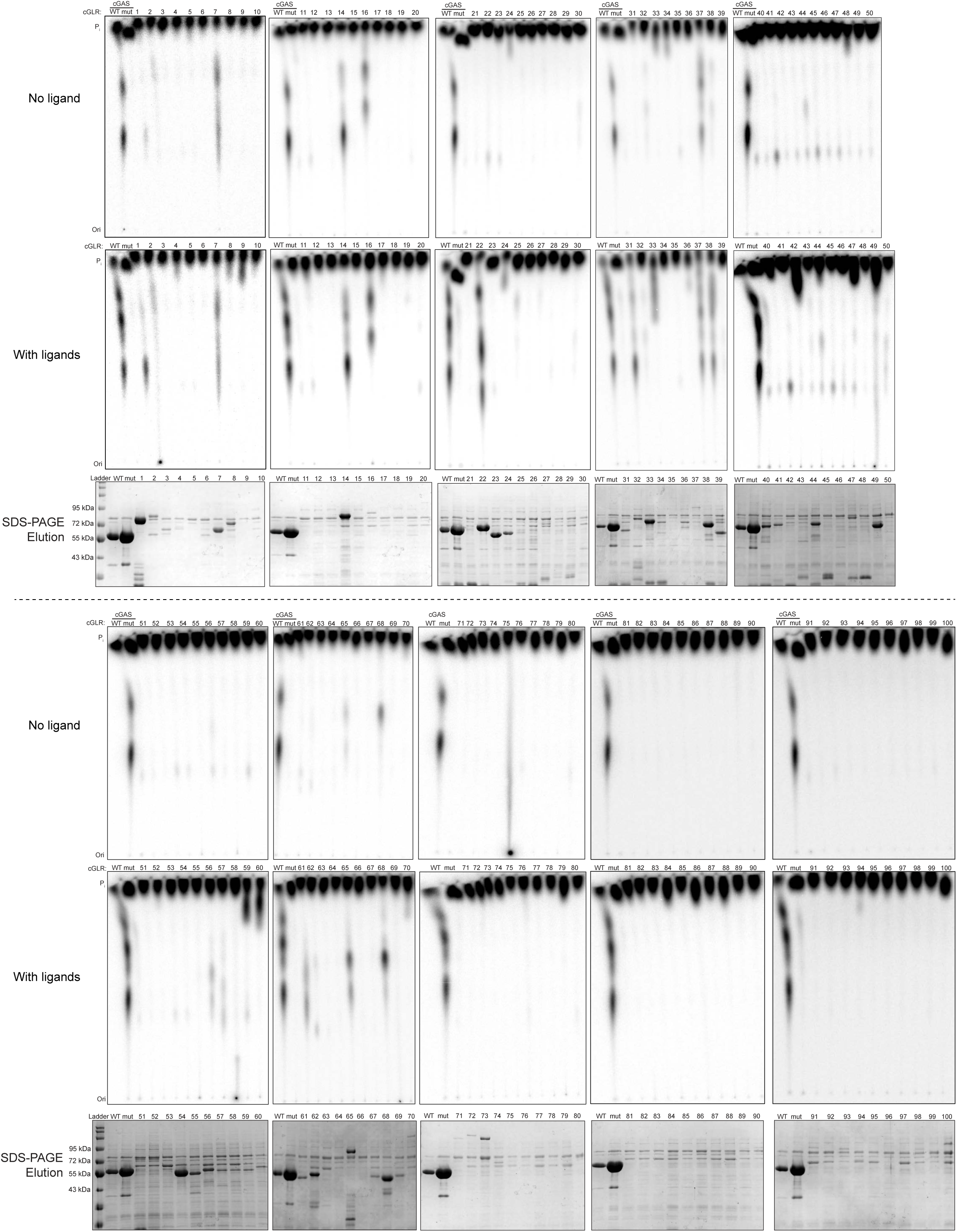
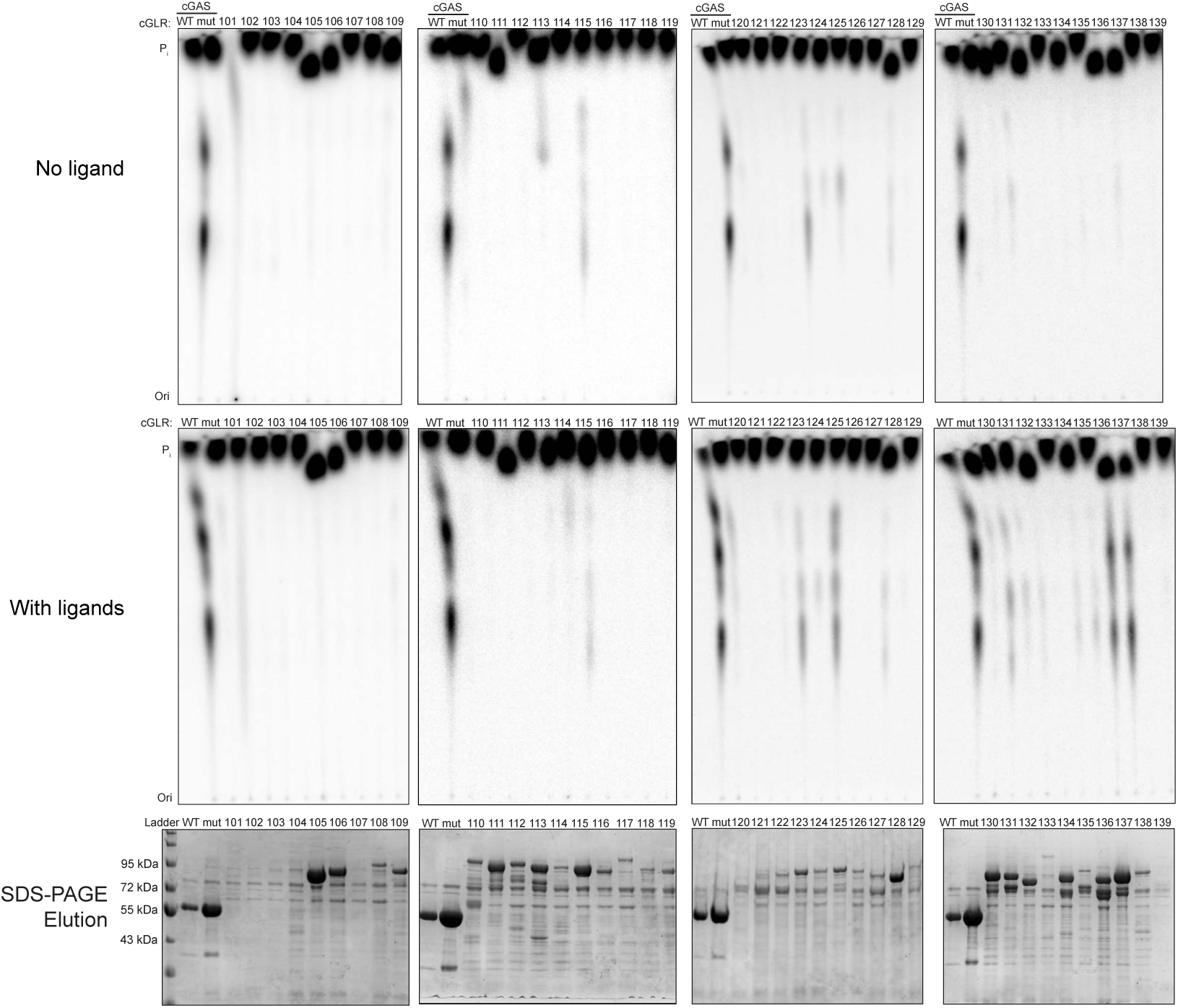
Biochemical screen of cGLRs from diverse animal species, related to. Figure 1 Primary data from a forward, biochemical screen of 140 animal cGLR proteins. Purified proteins were incubated with α^32^P-radiolabeled NTPs, and reaction products were visualized by PEI-cellulose TLC as in Figure 1D. Protein expression level and purity of each cGLR used in the screen are measured by SDS-PAGE and Coomassie stain analysis.

**Figure S4.**
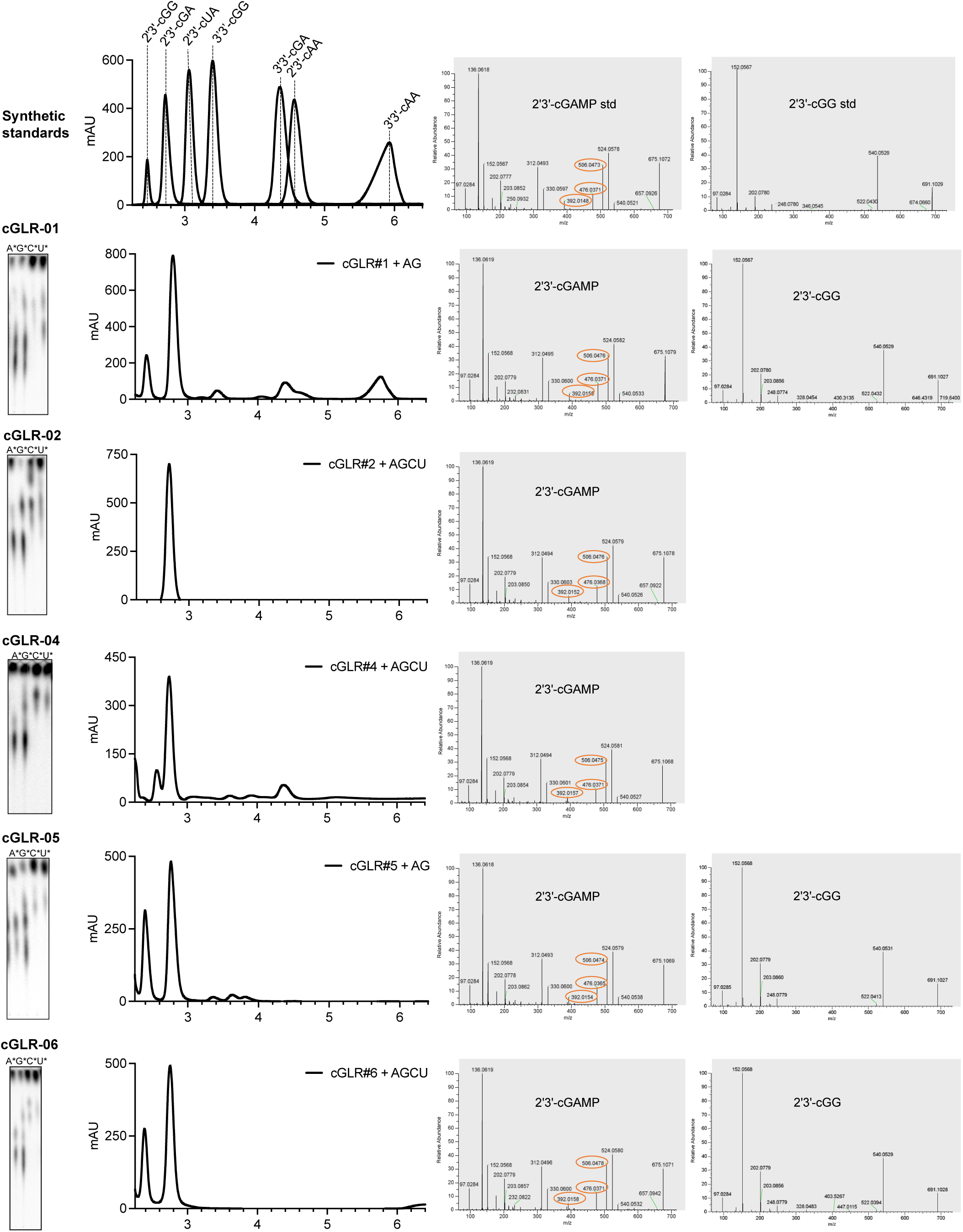
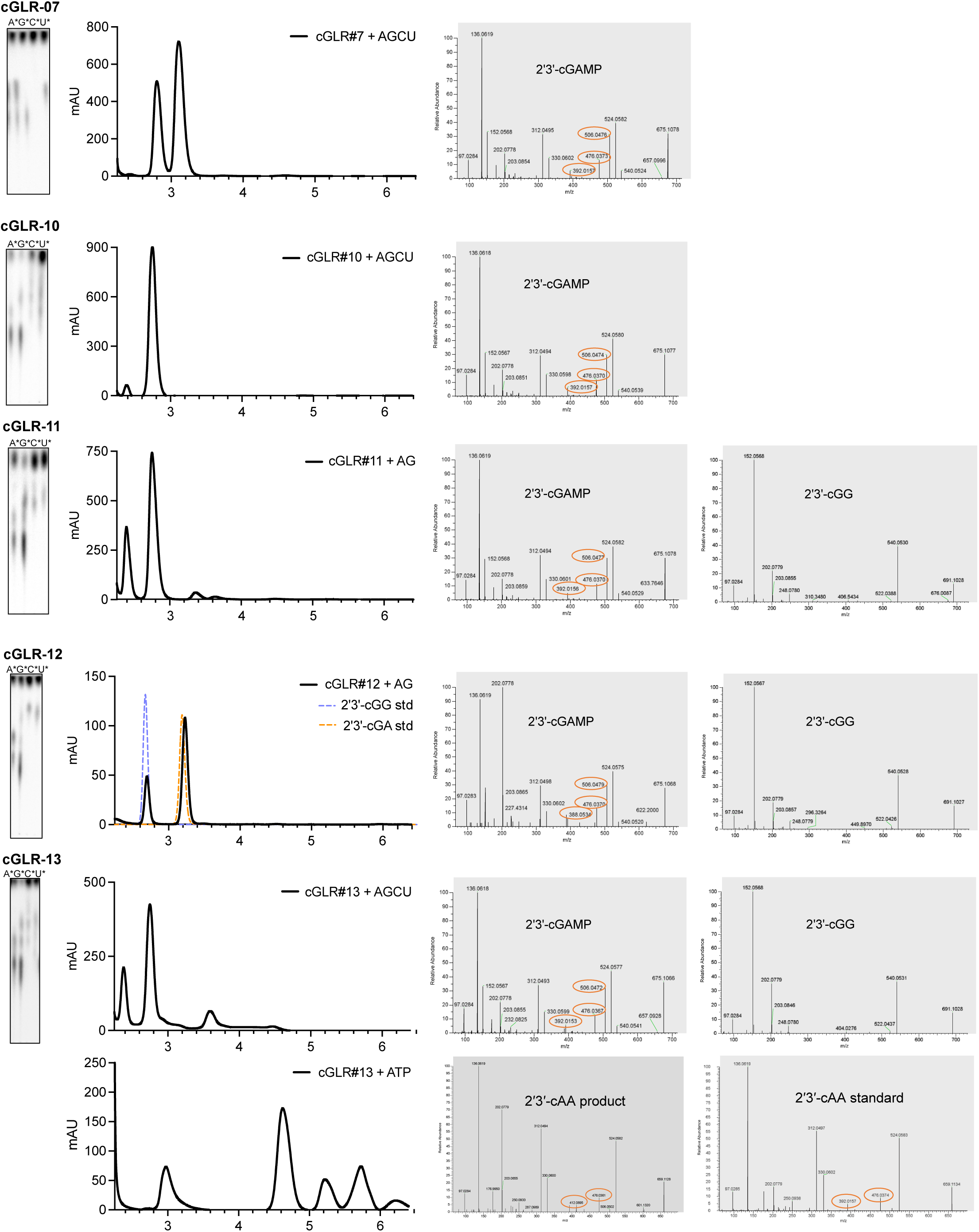
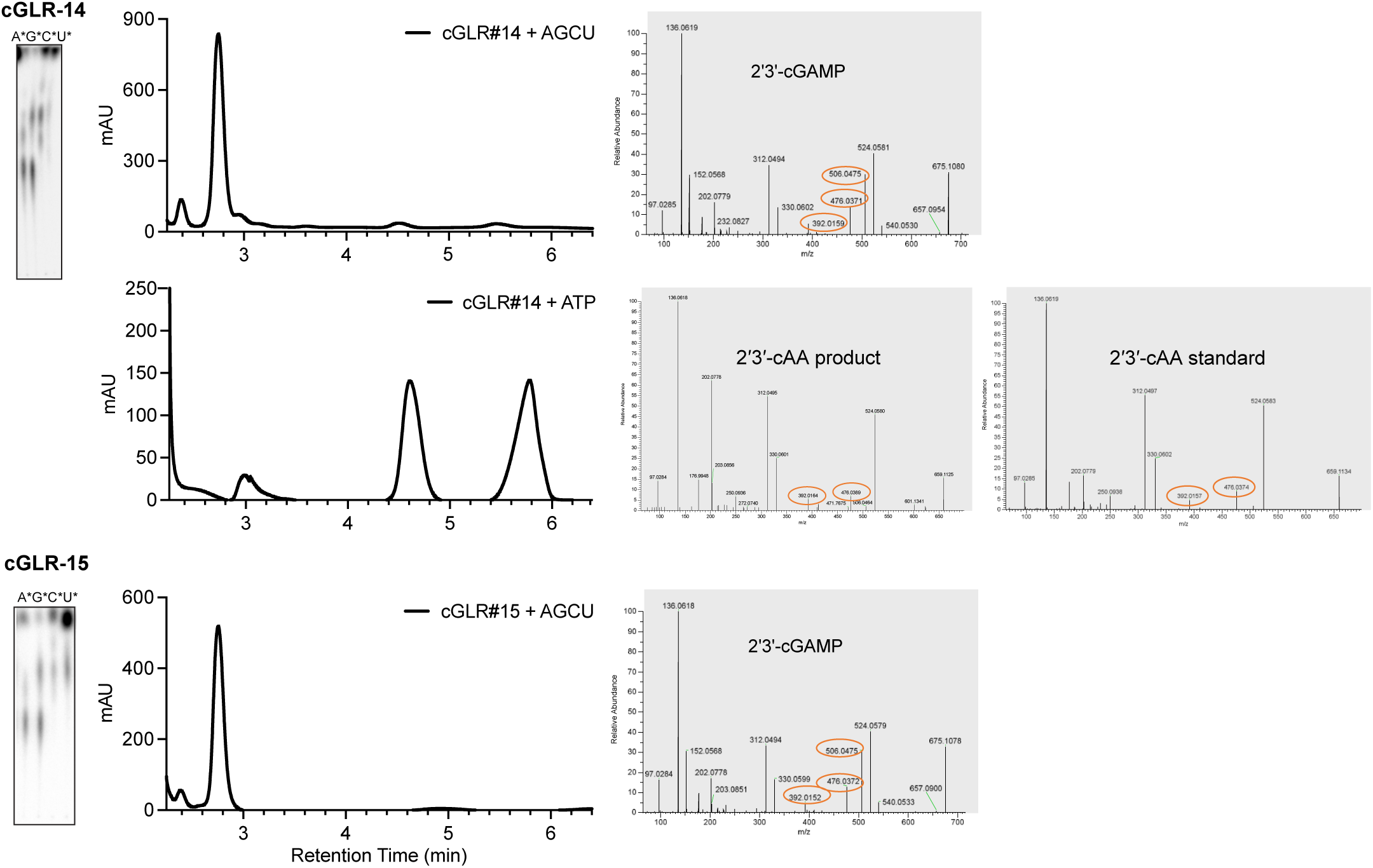
Identification of known cGLR nucleotide second messenger products, related to Figure 1. Combined biochemical deconvolution and LC-MS/MS analysis used to identify known cGLR nucleotide second messenger products. Active cGLR enzymes were incubated with unlabeled NTPs and each individual α^32^P-labeled NTP to reveal which nucleobases are incorporated into the major product species. Next, cGLR major nucleotide products were confirmed using HPLC and MS/MS analysis compared to synthetic standards of all previously known cyclic dinucleotide species.

**Figure S5.**
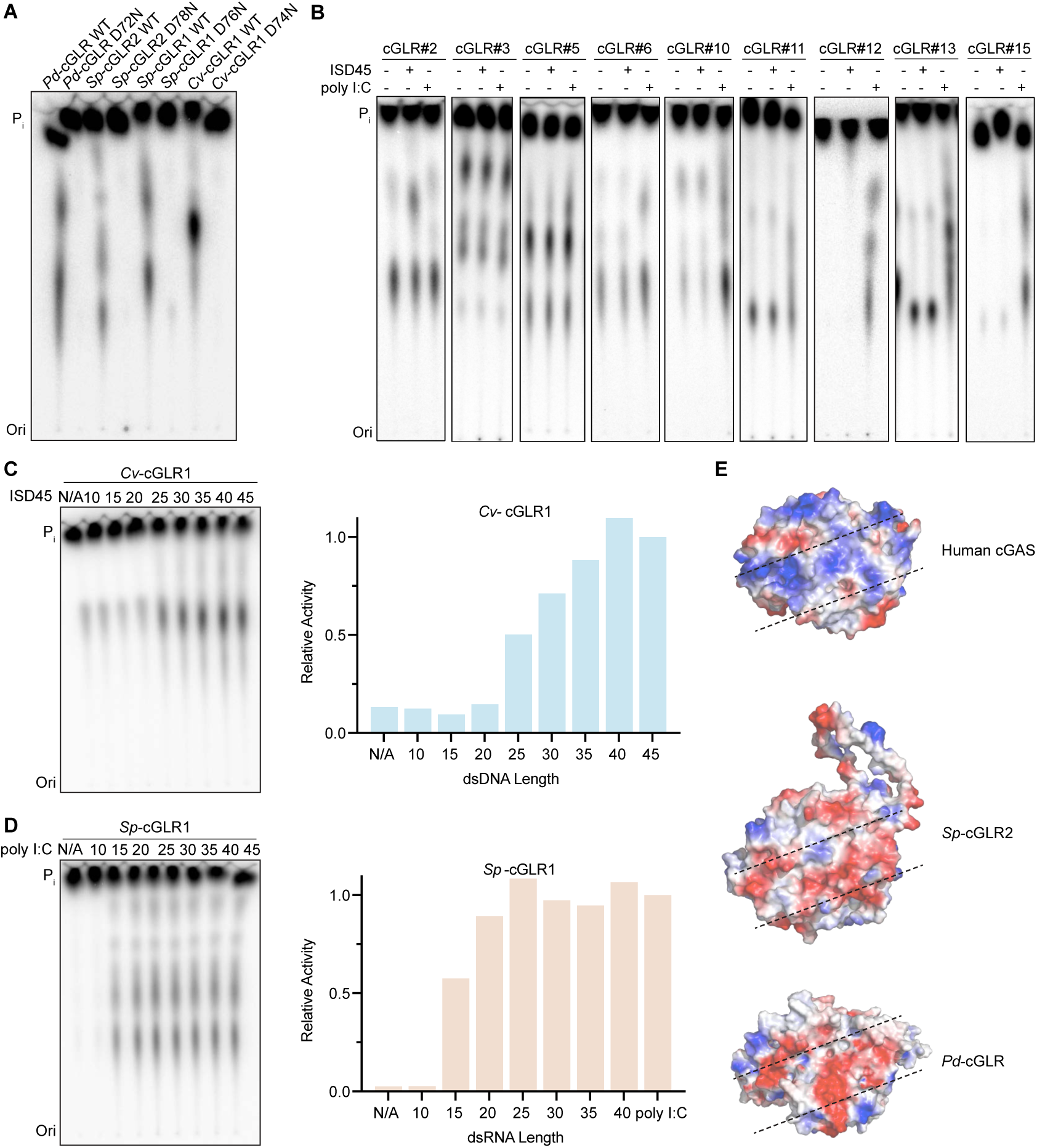
Analysis of cGLR activating ligand specificities, related to Figure 2. (A) Mutation to one of the key residues in the cGLR active site [DE]h [DE]h [X50–90] h[DE]h motif disrupts all enzymatic activity and confirms the specificity of metazoan cGLR nucleotide second messenger synthesis. Data are representative of n = 3 independent experiments. (B) Biochemical deconvolution of the activating ligand specificity of each active cGLR enzyme identified in the biochemical screen. cGLR enzymes are numbered according to Figure 1A. cGLR-01, -02, -03, -04, -05, -06 respond to an unknown ligand; cGLR-07, -08 respond to dsDNA; cGLR-09, -10, -11, -12, -13, -14, -15 respond to dsRNA. Data are representative of n = 3 independent experiments. (C,D) Thin layer chromatography analysis and quantification of enzyme activity of *Cv*-cGLR1 and *Sp*-cGLR1 in the presence of a panel of synthetic nucleic acid ligands. *Cv*-cGLR1 and *Sp*-cGLR1 respond to long double-stranded nucleic acid ligands. Data are representative of n = 3 independent experiments. (E) Surface charge of the ligand binding groove of human cGAS (PDB: 6CT9), *Pd*-cGLR (AlphaFold2 model) and *Sp*-cGLR2 (AlphaFold2 model).

**Figure S6.**
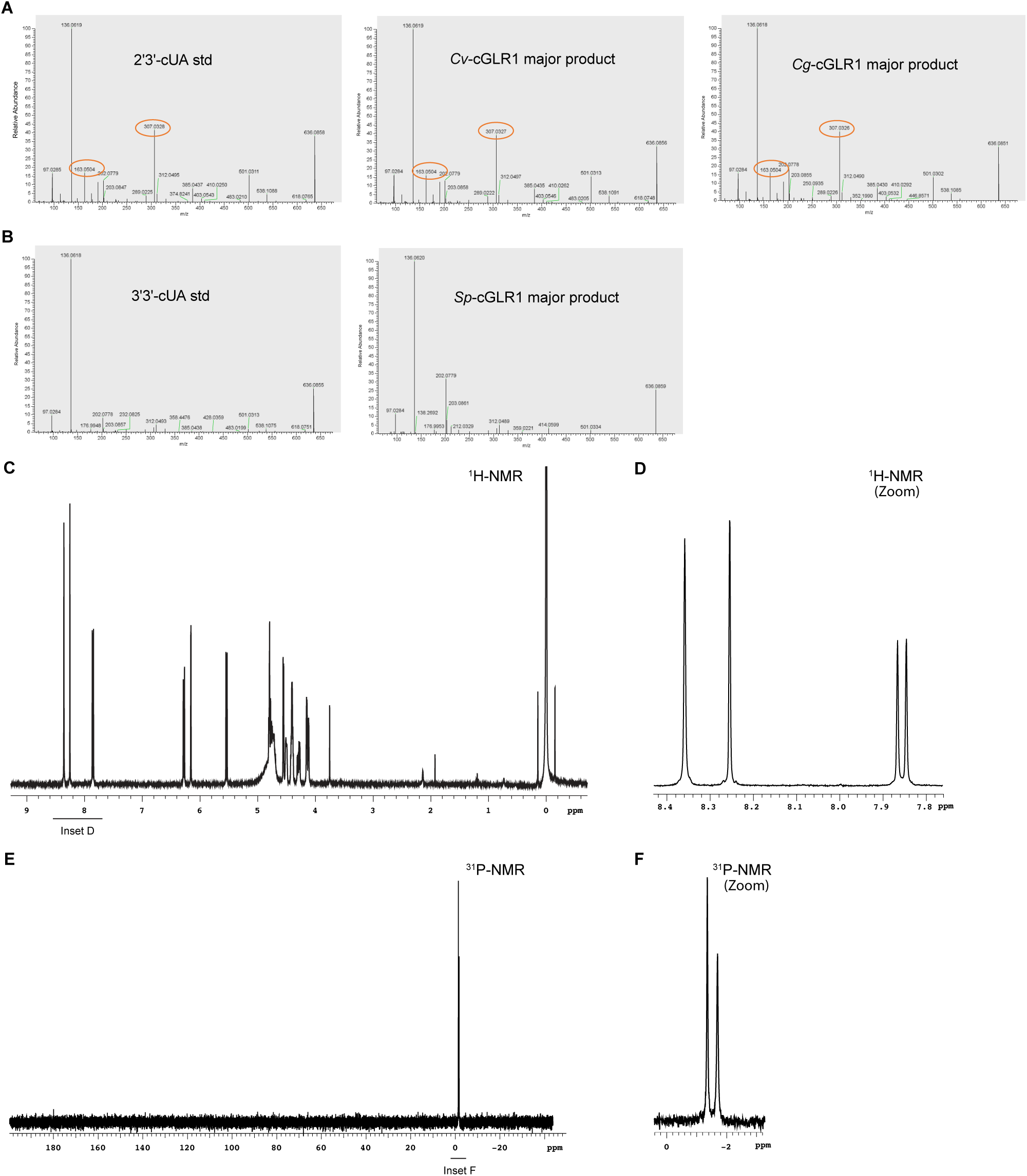
Identification of novel cGLR nucleotide second messenger products, related to Figure 3. Combined biochemical deconvolution and LC-MS/MS analysis used to identify novel cGLR nucleotide second messenger products using an approach similar to Figure S4. (A) MS/MS analysis following HPLC analysis shown in Figure 3 compared to synthetic standards confirms the major products of *Cv*-cGLR1, *Cg*-cGLR1 and (B) *Sp*-cGLR1 as 2′3′-cUA and 3′3′-cUA respectively. NMR analysis (C–F) further confirms the *Cv*-cGLR1 and *Cg*-cGLR1 major product 2′3′-cUA as a metazoan cyclic dinucleotide containing a pyrimidine base. (C–D) 2′3′-cUA proton-NMR spectrum (C) and associated magnified spectrum (D). ^1^H (400 MHz): δ_Η_ 8.36 (s, 1H), 8.26 (s, 1H), 7.85 (d, *J* = 8.2 Hz, 1H), 6.28 (d, *J* = 8.6 Hz, 1H), 6.16 (appt. d, *J* = 1.6 Hz, 1H), 5.54 (d, *J* = 8.2 Hz, 1H), 4.82–4.68 (m, 2H), 4.55 (d, *J* = 3.9 Hz, 1H) 4.52–4.38 (m, 1H), 4.37–4.22 (m, 4H), 4.19–4.10 (m, 2H). (E–F) 2′3′-cUA phosphate-NMR spectrum (E) and associated magnified spectrum (F). ^31^P{^1^H} NMR (162 MHz): δ_P_ −1.36 (s, 1P), −1.69 (s, 1P).

**Figure S7.**
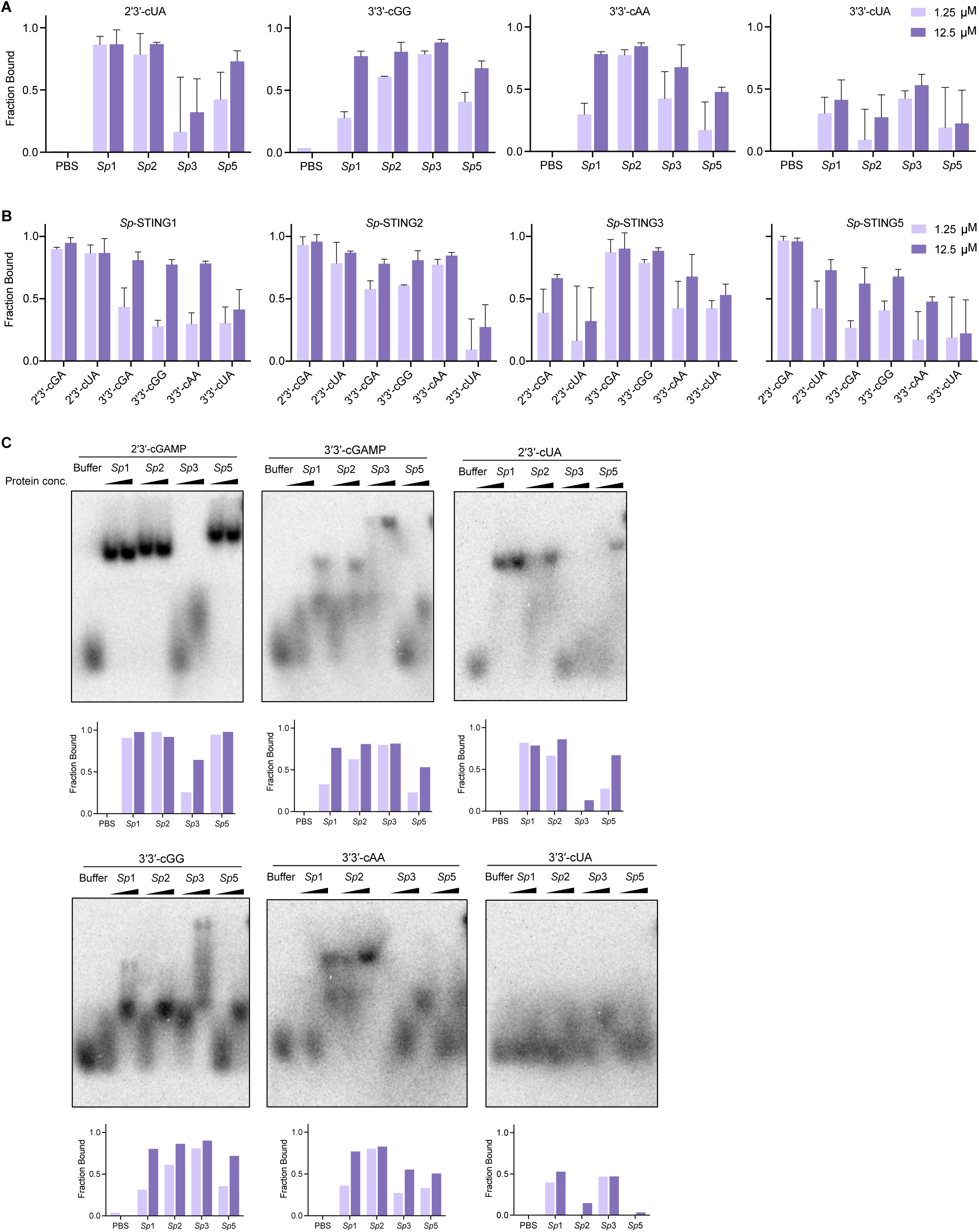
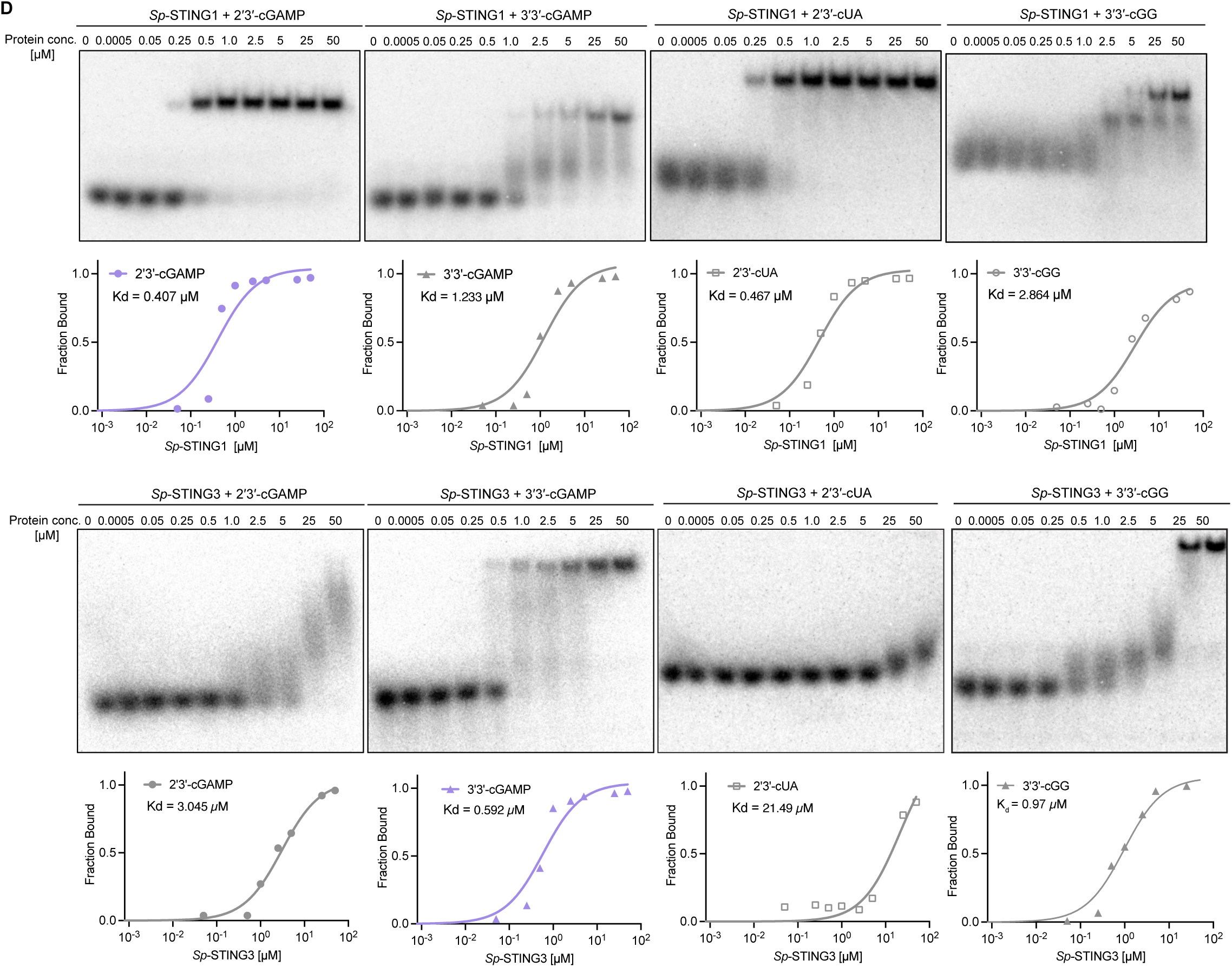
Biochemical analysis of *S. pistillata* STING cyclic dinucleotide recognition specificity, related to Figure 4. (A,B) Quantification of EMSA analysis of the binding of *Sp*-STING1, *Sp*-STING2, *Sp*-STING3 with 2′3′-cUA, 3′3′-cGG, 3′3′-cAA and 3′3′-cUA. Data are the mean ± std of n = 2 independent experiments. (C) Primary EMSA analysis data and quantification of the binding of *Sp*-STING1, *Sp*-STING2, *Sp*-STING3 with 2′3′-cGAMP, 3′3′-cGAMP, 2′3′-cUA, 3′3′-cGG, 3′3′-cAA and 3′3′-cUA. Data are representative of n = 2 independent experiments. (D) Primary EMSA analysis data and quantification of the binding affinity of *Sp*-STING1 and *Sp*-STING3 with 2′3′-cGAMP, 2′3′-cUA, 3′3′-cGG and 3′3′-cGAMP. Data are representative of n = 2 independent experiments.

**Figure S8.**
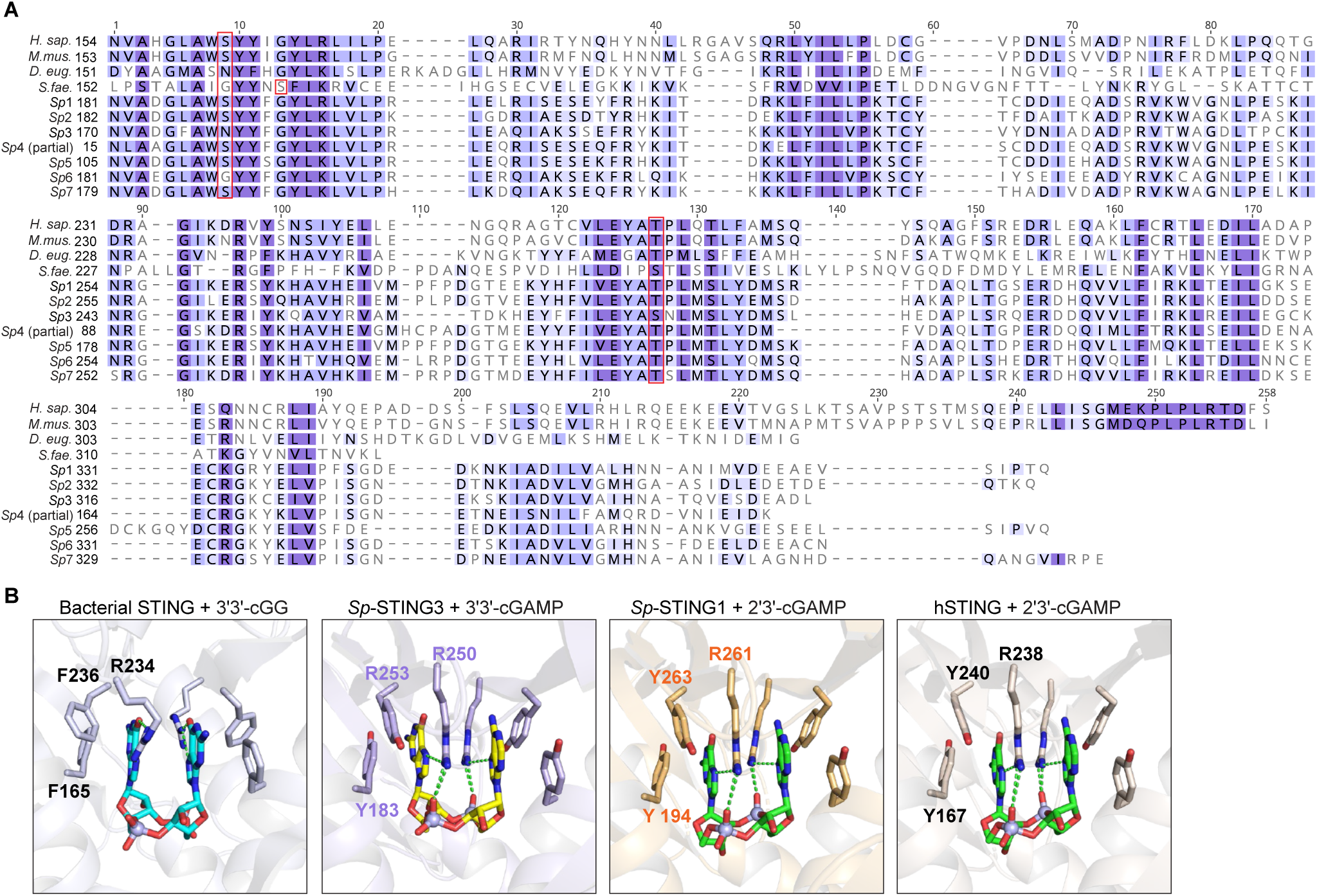
Sequence and structural analysis of *S. pistillata* STING receptors, related to Figure 5. (A) Sequence alignment of the cyclic dinucleotide binding domain (CBD) of *Sp*-STING proteins, STING from representative animal species, and the bacteria *S. faecium*. (B) Comparison of crystal and cryo-EM structures of the bacterial *Sf*-STING–3′3′-cGG complex (PDB: 7UN8), *Sp*-STING3–3′3′-cGAMP complex, *Sp*-STING1–2′3′-cGAMP complex and the human STING–2′3′-cGAMP complex (PDB: 4KSY) reveals conservation of specific cyclic dinucleotide contacts (Morehouse et al., 2020; Zhang et al., 2013).

## STAR METHODS

### RESOURCE AVAILABILITY

#### Lead Contact

Further information and requests for resources and reagents should be directed to and will be fulfilled by the Lead Contact, Philip Kranzusch.

#### Materials Availability

This study did not generate new unique reagents.

#### Data and Code Availability

Coordinates of the *Sp*-STING1–2′3′-cGAMP complex and *Sp*-STING3–3′3′-cGAMP structures have been deposited in the Protein Data Bank under the following accession numbers: 8EFM and 8EFN.

## EXPERIMENTAL MODEL AND SUBJECT DETAILS

### METHOD DETAILS

#### Bioinformatics and tree construction

Building on previous analyses (Slavik et al., 2021; Whiteley et al., 2019; Burroughs et al., 2015), animal cGLRs were identified using human cGAS and *Drosophila* cGLR1 as a query protein to seed a position-specific iterative BLAST (PSI-BLAST) search of the NCBI non-redundant protein database. Five rounds of PSI-BLAST searches were performed with an E value cut-off of 0.005 for inclusion into the next search round, BLOSUM62 scoring matrix, gap costs settings existence 11 and extension 1, and using conditional compositional score matrix adjustment. Hits from each round of PSI-BLAST were aligned using MAFFT (automatically determined strategy) (Katoh et al., 2019) and cGLRs were selected to refine the position-specific score matrix (PSSM) for the next round of search based on the presence of a conserved nucleotidyltransferase domain with a h[QT]GS [X8–20] [DE]h [DE]h [X50–90] h[DE]h motif. For the final round of selection, vertebrate and invertebrate hits were clustered separately using MMSeq2 (Steinegger and Söding, 2017). Major clusters of cGAS, Mab21L1, -L2, -L3 and MB21D2 from vertebrate genomes were identified and included for tree construction based on annotation of sequences, while invertebrate metazoan proteins were manually aligned and analyzed as individual clusters. Manual analysis and curation of candidate cGLR sequences was performed based on MAFFT protein alignment (automatically determined strategy) and predictive structural homology using HHPred (Söding et al., 2005) and AlphaFold2 (Jumper et al., 2021). Sequences were selected based on predicted structural homology to cGAS and conservation of a G[S/G] activation loop motif and a [E/D]h[E/D] X50–90 [E/D] catalytic triad.

To generate the phylogenetic tree in Figure 1A, cGLR sequences identified from PSI-BLAST were aligned using MAFFT (FFT-NS-i iterative refinement method) and truncated to the cGLR domain based on alignment with human cGAS. MMSeq2 was used to remove protein redundancies (minimum sequence identity = 0.95, minimum alignment coverage = 1) and the final aligned 3,138 sequences were used to construct a phylogenetic tree in Geneious Prime (v2022.1.1) using FastTree with no outgroup. iTOL was used for tree visualization and annotation (Letunic and Bork, 2021). Taxonomic and conserved domain analyses were performed using metadata associated with each nonredundant protein record in NCBI. Isoelectric point of cGLR proteins and bacterial CD-NTases were calculated using the DTASelect algorithm (Kozlowski, 2016). Full data for cGLR sequences and taxonomy distribution are included in Table S1. Information on the conserved domains fused to cGLR proteins are included in Table S2 and calculated isoelectric point of cGLR and CD-NTases are included in Table S4.

To generate the bubble plot in Figure 4B, animal STING sequences were identified using PSI-BLAST search with the same parameters as above. Manual analysis of identified cGLR and STING sequences was performed to determine the number of cGLR and STING genes encoded in each animal genome. Taxonomic analysis was performed using metadata associated with each organism in NCBI. Full data of for the number of cGLR and STING genes in each species as well as the associated taxonomy distribution are included in Table S5. PROMALS3D was used for Structure guided alignment of human STING (PDB 4KSY), *Drosophila* STING (PDB 7MWY), bacterial STING (PDB 7UN8), and seven coral STING proteins in Figure 5B and Supporting Figure S8A was prepared with PROMALS3D (Pei and Grishin, 2014) and visualized and annotated with Geneious Prime.

#### Cloning and plasmid construction

Cloning and plasmid construction was performed as previously described (Whiteley et al., 2019). Briefly, most *cGLR* genes and all the coral *STING* genes were synthesized as gBlocks (Integrated DNA Technologies) with ≥18 base pairs of homology flanking the insert sequence and cloned into a custom pETSUMO2 or pETMBP vector by Gibson assembly (Zhou et al., 2018; Whiteley et al., 2019). All synthesized sequences are presented in Table S3. Plasmids were transformed into the *E. coli* strain Top10 (Invitrogen).

#### Protein expression and purification

For the cGLR biochemical screen, each enzyme was expressed as a recombinant protein with 6×His-SUMO2- or 6×His-MBP-fusion tag and purified from *E. coli* in a small-scale format as previously developed for bacterial CD-NTase proteins (Whiteley et al., 2019). Briefly, plasmids were transformed into *E. coli* strain BL21-RIL (Agilent) and grown as overnight cultures in ∼3 mL of MDG media at 37°C with 230 RPM shaking. Overnight cultures were used to inoculate ∼10 mL M9ZB cultures grown at 37°C until OD_600_ reached 2.5–3.0, then induced with 0.5 mM IPTG and incubated overnight at 16°C with 230 RPM shaking. Bacterial pellets were resuspended and sonicated in lysis buffer (20 mM HEPES-KOH pH 7.5, 400 mM NaCl, 30 mM imidazole, 10% glycerol and 1 mM DTT), clarified by centrifugation at 3,200 × g (Eppendorf 5810R centrifuge) for 15 min at 4°C, and then proteins were purified from the supernatant by Ni-NTA purification (Qiagen) using 100 μL of packed resin and a spin column format. Recombinant protein was eluted in elution buffer (20 mM HEPES-KOH pH 7.5, 400 mM NaCl, 300 mM imidazole, 10% glycerol and 1 mM DTT) and buffer exchanged into assay buffer (20 mM HEPES-KOH pH 7.5, 250 mM KCl, 1 mM TCEP and 10% glycerol) using 10-kDa molecular weight cut-off spin column (Amicon). Recombinant proteins were used immediately for *in vitro* nucleotide synthesis reactions and analyzed by SDS–PAGE followed by Coomassie staining, shown in Figure S3.

Large-scale purification of recombinant cGLR and STING proteins was performed as previously described (Zhou et al., 2018; Slavik et al., 2021). Briefly, proteins were expressed as above in 2–4 1× liter M9ZB cultures, lysed in lysis buffer (20 mM HEPES-KOH pH 7.5, 400 mM NaCl, 30 mM imidazole, 10% glycerol and 1 mM DTT), and purified with Ni-NTA resin (Qiagen). Ni-NTA resin was washed with lysis buffer supplemented to 1 M NaCl and eluted with lysis buffer supplemented to 300 mM imidazole. Elution for cGLR-03, -04, -05, -10, -11, -15 proteins were buffer exchanged into storage buffer (20 mM HEPES-KOH pH 7.5, 250 mM KCl, 1 mM TCEP and 10% glycerol) and concentrated to >10 mg mL^−1^ before flash freezing with liquid nitrogen and storage at −80°C. cGLR-01, -02, -06, -07, -08, -09, -12, -13, -14 proteins were further purified by dialyzing into 20 mM HEPES-KOH pH 7.5, 150–300 mM NaCl, 1 mM DTT and 10% glycerol. For cGLR-07 and -08, the SUMO2 tag was removed with recombinant human SENP2 protease (D364–L589, M497A) during dialysis, and for cGLR -12, -13 and -14 the MBP tag was removed with recombinant TEV protease during dialysis. cGLR proteins were then purified by ion-exchange using a 5 mL HiTrap Heparin HP column (Cytiva) (cGLR-02, -06, -07, -08, -09, -12, -13, -14) or 5 mL Q column (Cytiva) (cGLR-01) and eluted with a gradient of NaCl from 150 mM to 1 M. Target protein fractions were pooled and further purified by size-exclusion chromatography using a 16/600 Superdex 75 column or 16/600 Superdex 200 column (Cytiva). Proteins were concentrated to >10 mg mL^−1^, flash frozen with liquid nitrogen, and stored at −80°C.

#### Biochemical screening of cGLR nucleotide second messenger synthesis activity

Biochemical screen of cGLR nucleotide synthesis activity was analyzed by thin layer chromatography (TLC) as previously described (Whiteley et al., 2019; Slavik et al., 2021). Briefly, 1 µL of purified cGLR protein was incubated with 50 μM of each unlabeled ATP/CTP/GTP/UTP and 0.5 μL α-^32^P-labeled NTPs (approximately 0.4 μCi each of ATP, CTP, GTP and UTP) in cGLR reaction buffer (50 mM Tris-HCl pH 7.5, 100 mM KCl, 10 mM MgCl_2_, 1 mM MnCl_2_ and 1 mM TCEP) at 37°C overnight. Reactions were supplemented with 1 μg poly I:C and 5 μM ISD45 dsDNA potential activating ligands as indicated. Mouse cGAS and its catalytically inactive mutant were used as controls (Zhou et al., 2018). Reactions were terminated with addition of 0.5 µL of Quick CIP phosphatase (New England Biolabs) to remove terminal phosphate groups from unreacted nucleotides. Each reaction was analyzed using TLC by spotting 0.5 µL on a 20 cm × 20 cm PEI-cellulose TLC plate (Millipore). The TLC plates were developed in 1.5 M KH_2_PO_4_ pH 3.8 until buffer was 1–3 cm from the top of plate and air-dried at room temperature and exposed to a phosphor-screen before imaging with a Typhoon Trio Variable Mode Imager (GE Healthcare).

To determine the activating ligand of cGLRs, 0.25–1 µM of each cGLR protein was incubated with 50 µM of each unlabeled NTPs and 0.5 μl α-^32^P-labeled NTPs in cGLR reaction buffer (50 mM Tris-HCl pH 7.5, 100 mM KCl, 10 mM MgCl_2_, 1 mM MnCl_2_ and 1 mM TCEP). Reactions were incubated either without any ligand or supplemented with 1 μg poly I:C or 5 μM ISD45 dsDNA as indicated at 37°C for 4 h or overnight.

To deconvolute the nucleotide composition of cGLR products, 0.25–1 µM of each cGLR protein was incubated with 50 µM of each unlabeled NTPs, 0.5 μL of individual α-^32^P-labeled NTP as indicated and necessary activating ligand determined in previous analysis. Reactions were terminated using Quick CIP and analyzed with TLC as described above. For P1 degradation assays, 4 µl of each reaction was treated with 0.5 µL Nuclease P1 (Sigma N8630) and 0.5 µL of Quick CIP in 1 × P1 digestion buffer (30 mM NaOAc pH 5.3, 5 mM ZnSO_4_, 50 mM NaCl) for 1 h at 37°C. Reactions were analyzed using TLC as described above.

#### Nucleotide purification and HPLC Analysis

All cGLR reactions were carried out at 37°C in standard reaction conditions as defined above: 50 mM HEPES-KOH pH 7.5, 50–100 mM KCl, 10 mM MgCl_2_, 1 mM MnCl_2_, 1 mM TCEP. Reactions containing 5 µM cGLR protein and necessary activating ligand according to Figure 2B were incubated overnight with 50 or 100 µM of each NTP as indicated. Samples were then spun through a 10-kDa molecular weight cut off spin column (Amicon) to remove protein and high molecular weight ligand. HPLC analysis was carried out as previously described (Slavik et al., 2021) at 40°C using a C18 column (Agilent Zorbax Bonus-RP 4.6×150 mm, 3.5-micron) with a mobile phase of 50 mM NaH_2_PO_4_ (pH 6.8 with NaOH) supplemented with 3% acetonitrile and run at 1 mL min^−1^. Nucleotide products of cGLRs were collected based on retention time using the fraction collector of the HPLC instrument (Agilent 1200 series) and concentrated using a speed vac before mass spectrometry analysis.

To prepare *Cg-*cGLR and *Cv*-cGLR1 nucleotide product for NMR analysis, 5 µM of enzyme was incubated with 1 mM UTP and ATP in the same reaction conditions indicated above. The reaction mixture was treated with Quick CIP for 3 h and heated for 1 h at 65°C before centrifugation at 4°C, 3,200 × g for 10 min and filtration through a 0.22 µm filter to remove precipitated protein. Nucleotide products were further purified by ion exchange using a HiTrap Q column and eluted in a 0–100% gradient of 1 M ammonium bicarbonate in water. Eluted fractions were concentrated by speedvac and resuspended in 500 µL of water and desalted by size exclusion chromatography with a Superdex 30 Increase 10/300 GL column (Cytiva) run in water.

#### Liquid chromatography-tandem mass spectrometry (LC-MS/MS) analysis

LC-MS/MS analysis samples were analyzed by the commercial company MS-Omics. The analysis was carried out using a UPLC system (Vanquish, Thermo Fisher Scientific) coupled with a high-resolution quadrupole-orbitrap mass spectrometer (Orbitrap Exploris 480 MS, Thermo Fisher Scientific) using an electrospray ionization interface operated in positive ionization mode. The UPLC was performed using a slightly modified version of the protocol described (Hsiao et al., 2018). Data were manually inspected to generate MS/MS spectra using Freestyle 1.4 (Thermo Fisher Scientific).

#### Chemical synthesis of cyclic dinucleotide standards

Synthetic nucleotide standards used for HPLC and mass spectrometry analysis were purchased from Biolog Life Science Institute: 3′3′-cGAMP (cat no. C 117), 2′3′-cGAMP (cat no. C 161), 3′2′-cGAMP (cat no. C 238), 2′3′-c-di-AMP (cat no. C 187), 2′3′-c-di-GMP (cat no. C 182), 3′3′-cUA (cat no. C 357).

Chemical synthesis of cyclic (uridine-(2′ −> 5′)-monophosphate-adenosine-(3′ −> 5′)-monophosphate) (2′3′-cUA / c[U(2′,5′)pA(3′,5′)p] / 2′3′-cUAMP / 2′–5′ / 3′–5′ cyclic UMP–AMP), sodium salt was performed as follows: 5 mmol of cyanoethyl phosphoramidite 5′-DMTr-2′-TBDMS-3′-CEP-N6-Bz-adenosine (ChemGenes, Wilmington, MA, USA, Cat. No. ANP-5671) were used as starting material for the synthesis of the corresponding phosphonate precursor with a standard oligonucleotide coupling protocol, originally developed for the synthesis of 3′3′-c-diGMP (Gaffney et al., 2010). 7 mmol (1.4 eq.) 3′-tBDSilyl-Uridine 2′-CED phosphoramidite (ChemGenes, Wilmington, MA, USA, Cat. No. ANP-5684) were used to form the protected dimeric linear precursor 5′-OH-3′-TBDMS-uridine-(2′→5′)-cyanoethyl-phosphate-2′-TBDMS-3′-H-phosphonate-N6-Bz-adenosine, followed by cyclization and removal of protection groups according to Gaffney et al 2010. Volatile components of the reaction mixture of 2′3′-cUA were evaporated under reduced pressure and stored at −70°C until further operations. 250 mL water were added, and the resulting suspension was placed in an ultrasonic bath at room temperature for 15 min, followed by 3 extraction cycles with 200 mL chloroform each. The combined organic phases were re-extracted with 200 mL water and the combined product-containing aqueous phase was filtered with a 0.45 µm regenerated cellulose (RC) filter and partially concentrated under reduced pressure to remove traces of chloroform. The raw product solution was diluted with water to 1200 mL and applied to a Q Sepharose Fast Flow anion exchange column (40–165 µm; 380 × 50 mm) Cl--form, previously regenerated with 2 M sodium chloride and washed with water. The column was washed with water (500 mL), followed by a gradient of 0–1 M triethylammonium bicarbonate buffer (TEAB, pH 7, 6000 mL) in water (detection wavelength 254 nm). The title compound eluted with ∼500 mM TEAB. Product-containing fractions were carefully concentrated to a final volume of approximately 400 mL with a rotary evaporator equipped with a drop catcher in-vacuo (Caution: Foaming due to gas evolution!). Subsequent purification of 2′3′-cUA was accomplished by preparative reversed phase medium pressure liquid chromatography (MPLC). The product solution was applied to a Merck LiChroprep®RP-18 column (15–25 µm; 435 × 50 mm) previously equilibrated with 100 mM triethylammonium formate (TEAF, pH 6.8) in water. Elution was performed with 100 mM TEAF, followed by a step-gradient of 1% and 3% 2-propanol, 20 mM TEAF (pH 6.8) in water.

For desalting, 2′3′-cUA fractions of sufficient purity (>99% HPLC) were applied to a YMC Triart Prep C18-S, 12 nm, S-15 column (15 µm; 470 × 50 mm), previously equilibrated with water. The column was washed with water to remove excess TEAF buffer. Afterwards, 2% 2-propanol in water was used to elute the desalted 2′3′-cUA. To generate the sodium salt form of 2′3′-cUA, pooled product-containing fractions were partially concentrated under reduced pressure and subsequently applied to a Toyopearl™ SP-650M cation exchange column (65 µm; 125 × 25 mm) Na+-form, previously regenerated with 2 M sodium chloride and washed with water. For elution the column was washed with water until no UV-absorbance was detectable at 254 nm anymore. After filtration and careful evaporation under reduced pressure, 519.16 µmol 2′3′-cUA, sodium salt, were isolated with a purity of 99.56 % HPLC (theoretical yield: 10.38%).

Formula (free acid): C_19_H_23_N_7_O_14_P_2_ (MW 635.38 g mol^−1^) UV-Vis (water pH 7.0): λ max 260 nm; ε 22500.

ESI-MS neg. mode: m/z 634 (M-H)-, m/z 656 (M-2H+Na)-.

#### NMR

All NMR analyses were conducted on a Varian 400-MR spectrometer (9.4 T, 400 MHz) as previously described (Whiteley et al., 2019). Samples were prepared by resuspending evaporated nucleotide samples in 500 µL D2O supplemented with 0.75% TMSP (3-(trimethylsilyl)propionic-2,2,3,3-d4) at 27°C. VnmrJ software (version 2.2C) was used to process data and generate figures. ^1^H and ^31^P chemical shifts are reported in parts per million (p.p.m.). J coupling constants are reported in units of frequency (Porritt and Hertzog) with multiplicities listed as s (singlet), d (doublet) and m (multiplet). These data appear in the figure legends of each NMR spectrum.

#### Electrophoretic mobility shift assay

Electrophoretic mobility shift assays were used to monitor the interactions between STING proteins and cyclic dinucleotides as previously described (Morehouse et al., 2020). Briefly, 20 nM of each α-^32^P labeled cyclic dinucleotide was incubated with STING proteins at indicated concentrations or with serial dilutions of STING protein ranging from 0.5 nM to 50 µM in a buffer containing 5 mM magnesium acetate, 50 mM Tris-HCl pH 7.5, 50 mM KCl, and 1 mM TCEP. Reactions were incubated at room temperature for 20 min before resolving on a 7.2-cm 6% nondenaturing polyacrylamide gel run at 100 V for 45 min in 0.5× TBE buffer. The gel was fixed for 15 min in a solution of 40% ethanol and 10% acetic acid before drying at 80°C for 45 min. The dried gel was exposed to a phosphor-screen and imaged on a Typhoon Trio Variable Mode Imager (GE Healthcare). Signal intensity was quantified using ImageQuant 5.2 software.

#### Crystallization and structure determination

Crystals of the *Sp*-STING1–2′3′-cGAMP complex and *Sp*-STING3–3′3′-cGAMP complex were grown at 18°C for 3–30 days using hanging-drop vapor diffusion. Purified STING proteins were diluted to 5 mg mL^−1^ in a buffer containing 20 mM HEPES-KOH pH 7.5, 75 mM KCl, 1 mM TCEP and incubated with 0.5 mM of cyclic dinucleotide as indicated. The mixture was incubated on ice for 10 min before used to set 96-well trays with 70 µL for each reservoir solution by mixing 200 nL of protein mixture and 200 nL of reservoir solution for each drop using Mosquito (SPT Labtech). Further optimized crystals for the *Sp*-STING1–2′3′-cGAMP complex were grown in EasyXtal 15-well trays (NeXtal Biotechnologies) with 400 μL reservoir solution and 2 μL drops set with a ratio of 1 μL of protein solution and 1 µL of reservoir solution. Optimized crystallization conditions were as follows: *Sp*-STING1–2′3′-cGAMP complex, 28% PEG 5000 MME, 100 mM Tris-HCl pH 8.6, 200 mM lithium sulfate; *Sp*-STING3–3′3′-cGAMP complex, 19% PEG 3350, 100 mM Bis-Tris propane, pH 6.4, 250 mM MgCl_2_. All crystals were harvested using reservoir solution supplemented with 10–25% ethylene glycol using a nylon loop.

X-ray diffraction data were collected at the Advanced Photon Source beamlines 24-ID-C and 24-ID-E. Data were processed with XDS (Kabsch, 2010) and Aimless (Evans and Murshudov, 2013) using the SSRL autoxds script (A. Gonzales, SSRL, Stanford, CA, USA). Experimental phase information for *Sp*-STING1 protein was determined for *Sp*-STING1 by preparing selenomethionine-substituted protein as previously described (Eaglesham et al., 2019) and using data collected from selenomethionine-substituted crystals. Anomalous sites were identified, and an initial map was generated with AutoSol within PHENIX (Liebschner et al., 2019). Structural modelling was completed in Coot (Emsley and Cowtan, 2004) and refined with PHENIX. Following model completion, the structure of the *Sp*-STING1–2′3′-cGAMP complex was used for molecular replacement to determine initial phases for the *Sp*-STING3–3′3′-cGAMP complex. Final structures were refined to stereochemistry statistics as reported in Table S6.

#### Quantification and Statistical Analysis

Statistical details for each experiment can be found in the figure legends and outlined in the corresponding methods details section. Data are plotted with error bars representing the standard deviation (SD).

## Notes

### Competing Interest Statement

The authors have declared no competing interest.

