## Supplementary material for "cGLRs are a diverse family of pattern recognition receptors in animal innate immunity": Li et al Table S6

**Table S6. Crystallographic Statistics, Related to Figure 5**

| | *Sp*-STING1–2′3′-cGAMP (SeMet) | *Sp*-STING1–2′3′-cGAMP | *Sp*-STING3–3′3′-cGAMP |
|---|---|---|---|
| **Data Collection** | | | |
| Resolution (Å)[a] | 39.92–2.17 (2.24–2.17) | 39.03–2.14 (2.20–2.14) | 38.84–1.73 (1.76–1.73) |
| Wavelength (Å) | 0.97918 | 0.97918 | 0.97918 |
| Space group | $P\ 6_1\ 2\ 2$ | $P\ 6_1\ 2\ 2$ | $P\ 2_1\ 2_1\ 2_1$ |
| Unit cell: a, b, c (Å) | 89.80, 89.80, 174.83 | 90.13, 90.13, 174.40 | 51.47, 82.70, 84.80 |
| Unit cell: $\alpha$, $\beta$, $\gamma$ (°) | 90.0, 90.0, 120.0 | 90.0, 90.0, 120.0 | 90.0, 90.0, 90.0 |
| Molecules per ASU | 2 | 2 | 2 |
| Total reflections | 1574999 | 324475 | 472440 |
| Unique reflections | 22786 | 23858 | 38535 |
| Completeness (%)[a] | 100.0 (99.6) | 99.8 (98.1) | 99.9 (98.5) |
| Multiplicity[a] | 69.1 (46.1) | 13.6 (8.9) | 12.3 (3.6) |
| $I/\sigma I$[a] | 18.4 (4.1) | 14.6 (2.0) | 12.4 (1.7) |
| CC(1/2)[b] (%)[a] | 99.9 (93.1) | 99.8 (48.4) | 99.8 (67.8) |
| Rpim[c] (%)[a] | 3.5 (75.3) | 3.6 (78.9) | 3.6 (40.6) |
| Sites | 4 | | |
| **Refinement** | | | |
| Resolution (Å) | | 38.87–2.13 | 38.84–1.73 |
| Free reflections | | 2095 | 2005 |
| R-factor / R-free | | 19.7 / 22.7 | 17.1 / 19.5 |
| Bond distance (RMS Å) | | 0.002 | 0.004 |
| Bond angles (RMS °) | | 0.532 | 0.728 |
| **Structure/Stereochemistry** | | | |
| No. atoms: protein | | 2868 (2 copies) | 2852 (2 copies) |
| No. atoms: ligand | | 45 | 45 |
| No. atoms: solvent | | 196 | 223 |
| Average B-factor: protein | | 29.79 | 25.61 |
| Average B-factor: ligand | | 18.45 | 12.90 |
| Average B-factor: water | | 34.70 | 35.39 |
| Ramachandran plot: favored | | 97.60% | 98.27% |
| Ramachandran plot: allowed | | 2.40% | 1.73% |
| Ramachandran plot: outliers | | 0.00% | 0.00 |
| Rotamer outliers | | 0.96% | 0.98% |
| MolProbity[d] score | | 1.32 | 1.12 |
| Protein Data Bank ID | | 8EFM | 8EFN |

[a] Highest resolution shell values in parenthesis
[b] (Karplus and Diederichs, 2012)
[c] (Weiss, 2001)
[d] (Chen et al., 2010)
